# Evolutionary predisposition of NIN to function in nitrogen-fixing nodules

**DOI:** 10.1101/2025.08.21.671612

**Authors:** Jieyu Liu, Siqi Yan, Min Li, Defeng Shen, Michaela Tichá, René Bærentsen, Kasper Røjkjær Andersen, Floris Verbeek, Olga Kulikova, Rene Geurts, Ton Bisseling, Rik Huisman

**Author notes:** These authors contributed equally to this work: Jieyu Liu, Siqi Yan. These authors contributed equally to this work: Ton Bisseling, Rik Huisman. **Correspondence and requests for materials** should be addressed to René Geurts, Ton Bisseling or Rik Huisman.

## Abstract

Nitrogen-fixing nodule symbiosis is an ecologically and economically important trait in legumes and some related species. A critical step in the evolution of nodulation is the recruitment of NODULE INCEPTION (NIN); a homolog of the nitrate-sensing NIN-LIKE PROTEIN (NLP) transcription factors. However, whether adaptations have occurred in the NIN protein upon its recruitment in symbiosis remains elusive. Here we show that non-symbiotic NIN orthologs can function in intracellular infection and even nodule initiation, demonstrating that these properties of NIN predate the evolution of nodulation. Concurrent with the evolution of nodulation, symbiotic NIN proteins were optimized for their role in symbiosis by acquiring nitrate independent functionality, including constitutive nuclear localization. A single amino acid substitution in Arabidopsis AtNLP2 enhances its nuclear localization under low nitrate conditions, making it functionally comparable to a symbiotic NIN. These findings highlight that NIN was predisposed to function in nodulation at the time of its recruitment into nitrogen-fixing symbiosis. Our study provides novel insight in how and why non-symbiotic NIN orthologues could be recruited to function in nitrogen-fixing root nodule symbiosis.

## Introduction

Some plant species can establish a nitrogen-fixing root nodule endosymbiosis with diazotrophic rhizobium or *Frankia* bacteria. Nodulating species are members of the related orders Fabales, Fagales, Cucurbitales, and Rosales, known collectively as the Nitrogen-Fixing Clade (NFC)^1,2^. The evolution of nodulation depends on at least two critical events that occurred in the common ancestor of the NFC^3^. First, the recruitment of the common symbiotic signalling pathway from the more ancient arbuscular mycorrhizal endosymbiosis, which is widespread across the plant kingdom^4^. This signalling initiates both endosymbiotic interactions upon detecting a symbiotic microbe. Second, the recruitment of the transcription factor NODULE INCEPTION (NIN). In the common ancestor of the NFC, *NIN* gained a *cis*-regulatory element in its promoter, placing its expression directly under the control of the common symbiotic signalling pathway^5^. As the most upstream nodulation-specific transcription factor, NIN functions as the master regulator essential for multiple steps of nodule formation and functioning^6–13^. However, whether adaptations in the NIN protein itself were critical to commit to such a key function in nodulation remains elusive.

NIN is part of the NIN-LIKE PROTEIN (NLP) family^14^. NLPs are primary nitrate sensors, playing a central role in nitrate-induced signalling^15^. The *NIN* orthogroup arose upon a duplication in an ancestral dicot plant species, predating the evolution of the NFC^16^. Therefore, non-nodulating plant species outside the NFC have *NIN* orthologs. NIN orthologs from within- and outside the NFC induce distinct downstream genes. Symbiotic NINs induce genes involved in nodule development such as *NF-YA* and *NF-YB* encoding subunits of the NUCLEAR FACTOR Y complex^17^, while non-symbiotic NIN orthologs like *Arabidopsis thaliana* AtNLP2 regulate nitrate responsive genes^18^. Moreover, the subcellular localization of symbiotic and non-symbiotic NIN proteins is different. Symbiotic NINs are nuclear localized^10,19^, whereas non-symbiotic NINs are generally cytoplasmic under low nitrate conditions and only shuttle to the nucleus upon sensing high nitrate^18,20^. This suggests that changes also occurred in the NIN protein when it was recruited in nodulation.

### Non-symbiotic NIN orthologs are partially functional in nodulation

To test to what extent non-symbiotic NIN proteins can function in nodulation, we selected orthologs representing a range of phylogenetic distances to the legume experimental model Medicago (*Medicago truncatula*). These include *PanNIN* of the nodulating non-legume species Parasponia (*Parasponia andersonii*), and the non-symbiotic orthologs of three species outside the NFC; *MeNIN1* and *MeNIN2* of cassava (*Manihot esculenta*), *AtNLP1*, *AtNLP2*, and *AtNLP3* of Arabidopsis, and *SlNIN* of tomato (*Solanum lycopersicum*)^14,19,21^ (Fig. 1a). In addition, the Medicago *NIN* paralog *MtNLP1* was included. We used these NIN homologs to trans-complement the Medicago *Mtnin-1* knock-out mutant, which can neither form infection threads nor nodules^9^. The experiments were conducted under 0.5 mM nitrate conditions. As codon usage in *NIN* orthologs of the different plant species is similar (Supplementary Table 1), we used native coding sequences driven by the Medicago *MtNIN* promoter^22^. These trans-complementation experiments showed that nodules were efficiently formed on roots transformed with symbiotic *NIN* genes *MtNIN* and *PanNIN*. Notably, NIN orthologues from three non-NFC species; Arabidopsis (AtNLP2), cassava (MeNIN1) and tomato (SlNIN), were also able to restore infection thread formation (Fig. 1b,l-o). In contrast, *MtNLP1* was unable to restore infection thread formation, indicating a larger functional divergence in the NIN paralogue than in the orthologues. When trans-complementing *Mtnin-1* with *AtNLP2* also some infected nodule primordia were observed, whereas other non-symbiotic NINs and MtNLP1 did not induce such structures (Fig. 1b, Extended Data Fig. 1). These results show that all three non-symbiotic species have a NIN orthologue that can restore infection thread formation in *Mtnin-1*, while the other NIN orthologues of Arabidopsis and cassava have lost this ability, likely due to neo-functionalisation after lineage-specific duplication.

**Fig. 1.**
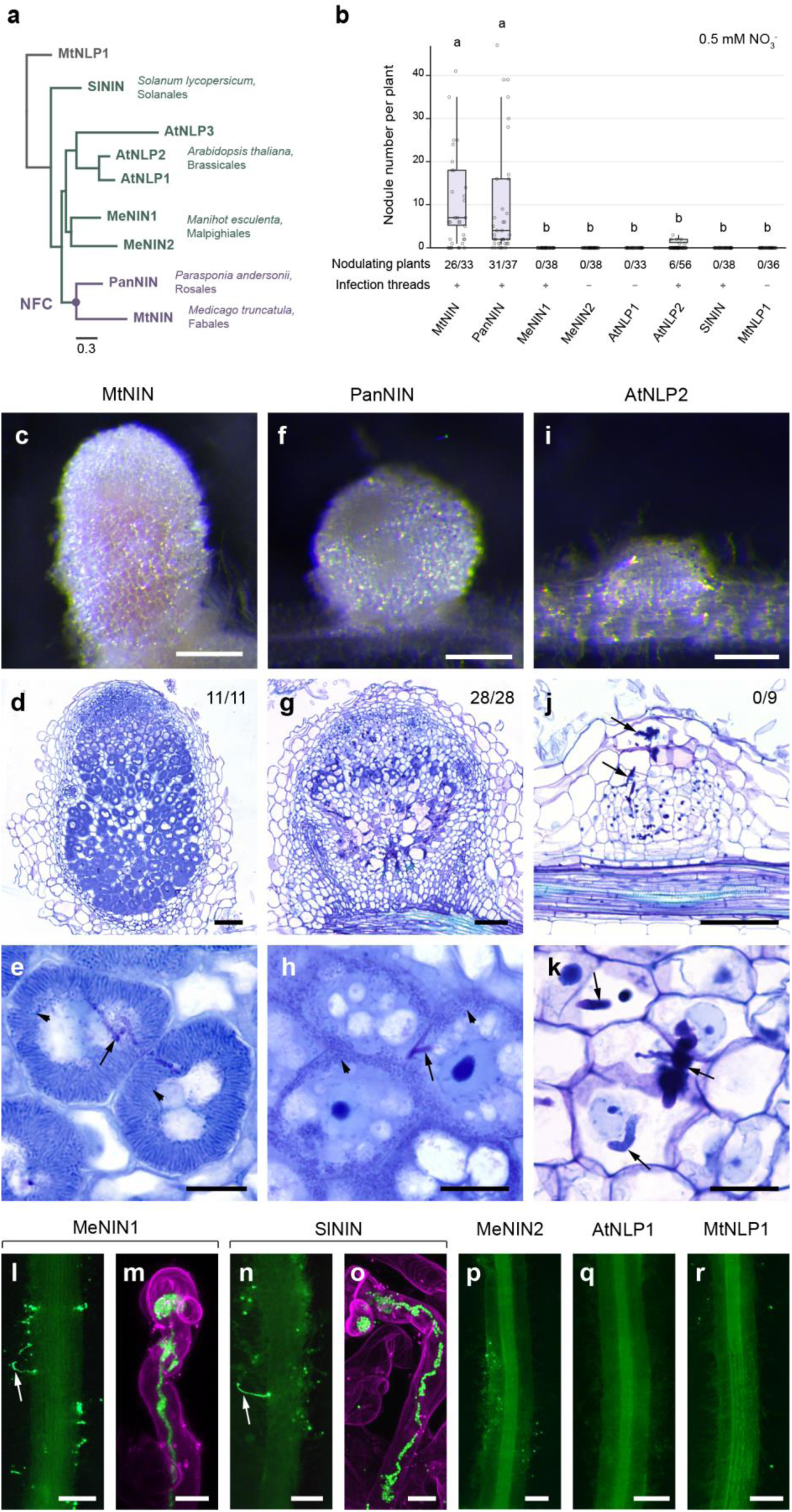
Non-symbiotic NIN orthologs are partially functional in nodulation. **a**, Phylogenetic tree of studied NIN orthologs and paralog MtNLP1. NFC, Nitrogen-Fixing Clade. Mt, *Medicago truncatula;* Pan, *Parasponia andersonii;* Me, *Manihot esculenta;* At, *Arabidopsis thaliana*; Sl, *Solanum lycopersicum.* **b**, Number of nodules formed on *Mtnin-1* mutant roots complemented with different NIN orthologs and a paralog. Plants were harvested at 4 weeks post inoculation with *Sinorhizobium meliloti* 2011 expressing *GFP*. Box plots show the number of nodules per nodulated plant. Lowercase letters indicate significant differences between samples (Kruskal-Wallis and post-hoc Dunn’s test, Benjamini-Yekutieli adjusted *p* < 0.05). Purple: symbiotic NIN, green: non-symbiotic NIN, grey: NIN paralog. **c**-**k** Images of nodules formed on *Mtnin-1* complemented with *MtNIN, PanNIN,* and *AtNLP2*. **c**, **f**, and **i**, Stereomicroscope images. Scale bars: 2 mm. **d**, **g**, and **j**, Longitudinal sections stained with toluidine blue. Numbers indicate nodules with released bacteria. Scale bars: 100 μm. **e**, **h**, and **k**, Magnification of nodule cells. Arrows indicate infection threads; arrowheads indicate released rhizobia. Scale bars: 20 μm. **l**, **n**, and **p-r**, Green fluorescence stereomicroscopy images showing infection threads (arrows) formed on the roots transformed with *MeNIN1* and *SlNIN*. Scale bars: 2 mm. **m**,**o**, Confocal images of transgenic roots stained with propidium iodide (magenta) showing infection threads (green) formed in the root hairs. Scale bars: 10 μm.

The nodules formed by native MtNIN were pink (Fig. 1c), indicating the accumulation of leghemoglobin, a hallmark of functional nitrogen-fixing nodules. Sections of these nodules showed that they have a wildtype zonation. The rhizobium bacteria in the cells of the fixation zone were fully differentiated/elongated like in wildtype nodules (Fig. 1d,e). In contrast, nodules formed on roots complemented with *PanNIN* were white. While bacteria were released into cells of these nodules, they did not fully differentiate, and premature senescence occurred in the basal part of the nodules (Fig. 1f-h). The Parasponia and Medicago lineages diverged early in the NFC^1^. The lack of full trans-complementation of *Mtnin-1* by *PanNIN* suggests that, after the recruitment of NIN by the common ancestor of the NFC, lineage-specific evolution still occurred. The nodule primordia formed on roots complemented with *AtNLP2* were arrested at an early stage of development (Fig. 1i-k). Although infection threads penetrated the nodule cells, rhizobia were not released.

To further investigate the evolution of NIN, we used ancestral sequence reconstruction to resurrect the NIN protein of the most recent non-symbiotic ancestor in the Rosid clade (NIN_Rosids_)^23,24^ (Supplementary Table 2). Although the presence of multiple non-conserved regions in NIN poses challenges for reliable ancestral sequence reconstruction, the structure of NIN_Rosids_ resembles that of extant NIN proteins (Extended Data Fig. 2). Introduction of NIN_Rosids_ in *Mtnin-1* mutant roots resulted in infection thread formation (Extended Data Fig. 2), supporting predisposition for symbiotic functioning of the ancestral Rosid NIN. We next asked whether the ability to initiate nodule primordia was already present in NIN of the NFC ancestor (NIN_NFC_). To test this, we resurrected NIN_NFC_ and used it to complement *Mtnin-1*. Besides infection threads, this resulted in rare but observable nodule primordia (Extended Data Fig. 2).

Taken together, these results show that *NIN* orthologs from non-nodulating species can partially function in nodulation, and the ability of NIN to induce nodule primordia was likely already present in the NFC ancestor, facilitating the evolution of nodulation.

### NIN recruitment in nodulation did not require major changes in the C-terminal DNA binding domain

Since non-symbiotic NIN proteins function with markedly lower efficiency compared to symbiotic NINs, changes likely occurred in NIN that contributed to its functioning in nodulation. As they are transcription factors, we first tested the ability of NIN orthologs to target symbiotic genes. In *Lotus japonicus,* the C-terminus of NIN, containing its DNA-binding domain, binds *cis*-regulatory elements of *LjNF-YA1* and *LjNF-YB1.* In contrast, the NIN paralog *LjNLP4* does not bind these sites ^25^. Thus, changes in the NIN C-terminus may enable it to regulate symbiotic genes. NIN-binding sites were identified in symbiotic NIN-targets *MtLBD16*, *MtCEP7*, *MtCLE13* as well as the Medicago orthologs of *LjNF-YA1* and *LjNF-YB1* (*MtNF-YA1*, *MtNF-YB16)*^26–28^ (Extended Data Fig. 3d). These sites resemble those bound by the non-symbiotic ortholog AtNLP2^18^. Using electrophoretic mobility shift assays (EMSA), we observed significant DNA mobility shifts after incubation with the C-terminus of MtNIN, PanNIN, MeNIN1, MeNIN2, AtNLP2 and SlNIN, indicating that these proteins can bind symbiotic NIN-binding sites *in vitro* (Fig. 2a, Extended Data Fig. 3a,b).

**Fig. 2.**
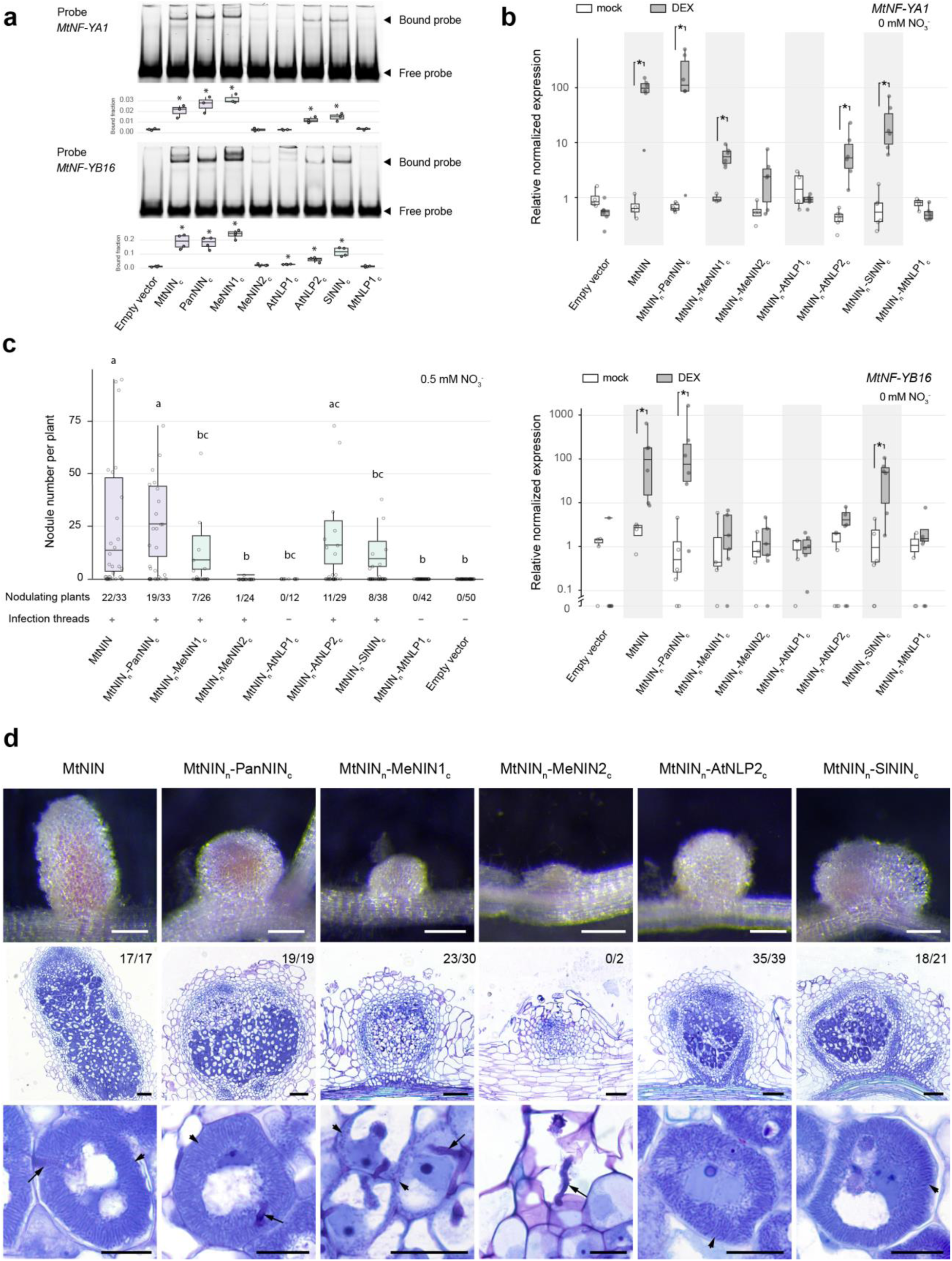
The C-terminal DNA binding domain of non-symbiotic NIN orthologs can function in symbiosis. **a**, Electrophoretic mobility shift assay (EMSA) testing the binding of the C-terminus of different NIN orthologs to NIN binding sites of *MtNF-YA1* and *MtNF-YB16*. Box plots show quantification of the bound fraction for each protein-probe combination (3-4 replicates). Asterisks indicate significant binding compared to empty vector control (Student’s t-test p < 0.05).**b**, qRT-PCR showing chimeric NIN proteins induce *MtNF-YA1* and *MtNF-YB16* expression in a transactivation assay. Medicago roots producing the N-terminus of MtNIN fused to the C-terminus of different NIN/NLPs and the rat glucocorticoid receptor were treated with dexamethasone (DEX) or DMSO (mock) for 16 hours. Expression levels were normalized to the average expression of mock treated empty vector roots. Asterisks indicate significant differences (Mann-Whitney *U-*test, p < 0.05). **c**, Complementation of Mt*nin-1* mutants with chimeric proteins, consisting of the N-terminus of MtNIN, fused to the C-terminus of different NIN/NLPs. Plants were harvested at 4 weeks post inoculation with *Sinorhizobium meliloti 2011* expressing *GFP*. Box plots show number of nodules per nodulated plant. Lowercase letters indicate significant differences between samples (Kruskal-Wallis and post-hoc Dunn’s test, Benjamini-Yekutieli adjusted *p* < 0.05). Purple: symbiotic NIN, green: non-symbiotic NIN. **d**, Images of nodules formed on *Mtnin-1* complemented with the chimeric proteins. Upper panels: stereomicroscope images; middle and bottom panels: longitudinal sections stained with toluidine blue. Numbers indicate nodules with released bacteria. Arrows: infection threads, arrowheads: released rhizobia. Scale bars: 2 mm (upper panels), 100 µm (middle panels) and 20 µm (bottom panels).

To test whether *in vitro* binding corresponds to *in vivo* ability to activate gene expression, we generated constructs in which the N-terminus of MtNIN was fused to the C-terminus of the tested NIN orthologs, and attached the glucocorticoid receptor (GR), allowing dexamethasone-controlled nuclear localization of the fusion proteins (Extended Data Fig. 4b). Medicago plants producing the fusion proteins were treated with dexamethasone for 16 hours. Following treatment, symbiotic NIN targets were induced by the symbiotic NINs as well as by chimers containing the non-symbiotic MeNIN1, MeNIN2, AtNLP2, and SlNIN C-termini (Fig. 2b, Extended Data Fig. 4a). The induction by the symbiotic NIN chimers was generally strong, while the induction by non-symbiotic chimers was variable, as they could induce some symbiotic NIN targets, but none of them significantly induced all targets. Similar results were obtained when using the full-length proteins (Extended Data Fig. 5).

To test whether the ability to induce symbiotic NIN targets is sufficient for functionality in symbiosis, we introduced chimeric NIN proteins (Extended Data Fig. 4c) into the *Mtnin-1* mutant roots. Nodules were formed on mutant plant roots complemented with constructs containing the C-terminus of symbiotic NINs and non-symbiotic NINs, MeNIN1, MeNIN2, AtNLP2, and SlNIN (Fig. 2c,d). On these roots, nodules were formed with phenotypes ranging from infected primordia (MeNIN2), release of rhizobia in nodule cells (MeNIN1), disorganized but differentiated rhizobia (AtNLP2), to radially organized differentiated rhizobia in pink nodules (SlNIN) (Fig. 2d). Neither nodules nor infection threads were formed on roots transformed with the empty vector control or constructs including the C-terminus of AtNLP1 or MtNLP1. Taken together, these results show that the C-terminus of most tested non-symbiotic NIN proteins allows activation of symbiotic targets, and is able to function in nodulation. This implies that the functionality of NIN in nodulation, including the release and differentiation of rhizobia, did not require major changes in its C-terminus.

### Nuclear localization is required for non-symbiotic NIN to function in a symbiotic context

Using the N-terminus from MtNIN enabled most non-symbiotic NINs to induce nodules, strongly suggesting that the major functional improvements for nodulation occurred in the N-terminus of NIN. Previous studies on the N-terminus of AtNLP7 showed that it has a nitrate sensing domain, and a phosphorylation site that is required for nitrate-triggered nuclear retention^29,30^. To test whether nitrate dependent nuclear localization was changed when NIN was recruited in nodulation, we determined the subcellular localization of NIN orthologs in Medicago nodules under 0 mM and 20 mM nitrate. This showed that symbiotic NINs were nuclear localized under both zero and high nitrate conditions. In contrast, AtNLP2, SlNIN, and MtNLP1 were localized in the cytoplasm and shuttled to the nucleus upon high nitrate application, consistent with previous studies ^18,31^. Similarly, MeNIN1 shuttled to the nucleus upon nitrate application, albeit with lower efficiency. The localization of AtNLP1 and MeNIN2 was nitrate independent; AtNLP1 remained in the cytoplasm, whereas MeNIN2 was nuclear localized in both conditions (Fig. 3a,b).

**Fig. 3.**
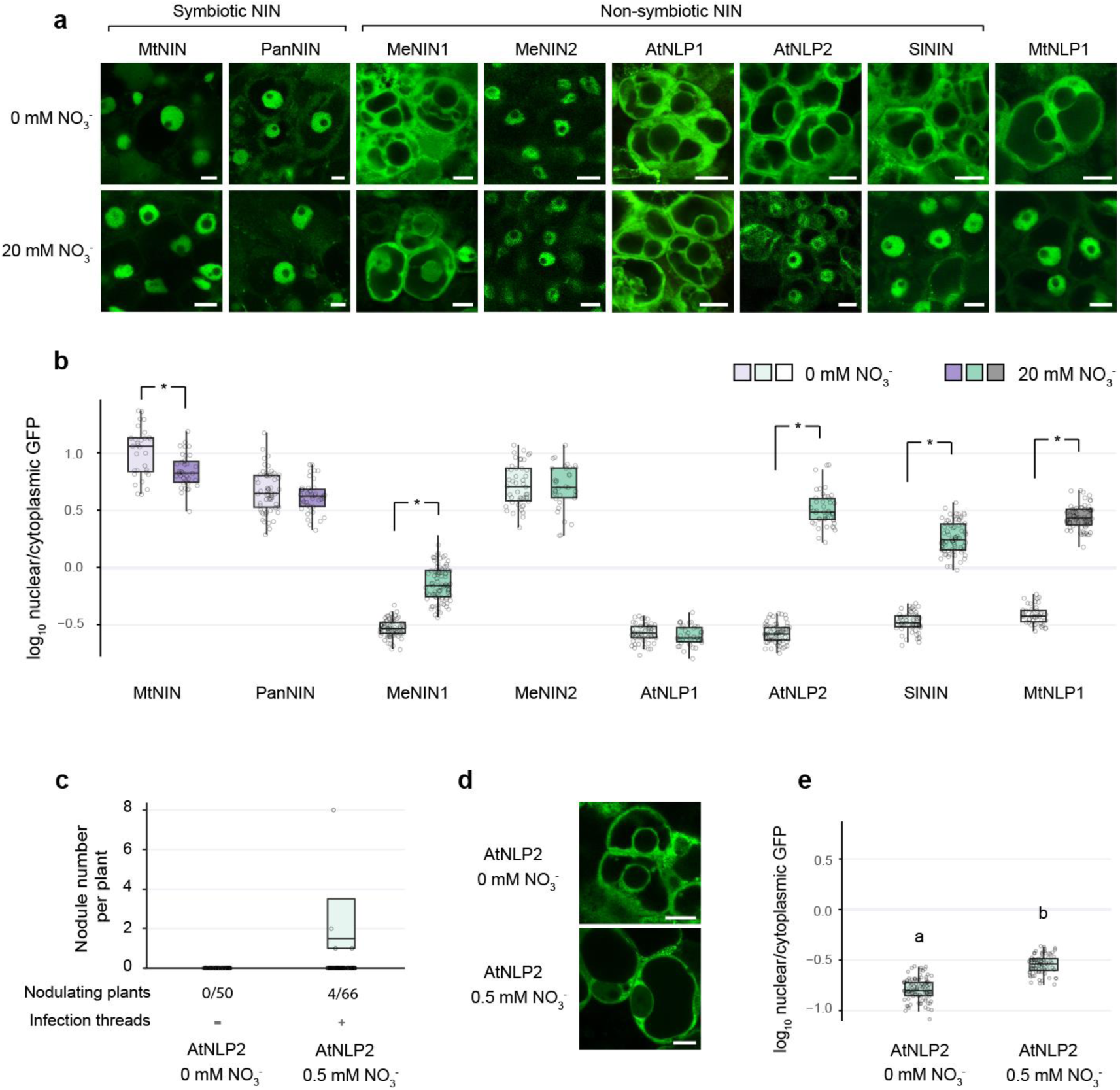
Nuclear localization of NIN orthologs is required for symbiotic function. **a**, Confocal images showing the subcellular localization of GFP-tagged *NIN* orthologs and a paralog in Medicago nodule infection zone cells at 0 mM and 20 mM nitrate. Scale bars: 10 µm. **b**, Quantification of subcellular localization shown in (**a**); Asterisks indicate significant differences (Mann-Whitney *U-*test, Benjamini-Yekutieli adjusted *p* < 0.05). **c**, Number of nodules formed on Mt*nin-1* mutant roots complemented with *AtNLP2* at 0 mM nitrate or 0.5 mM nitrate. Differences are not significant (Mann-Whitney U-test, *p* < 0.05). **d**, Confocal images showing subcellular localization of AtNLP2 under 0 mM or 0.5 mM nitrate. Scale bars: 10 μm. **e**, Quantification of subcellular localization in (**d**). Lowercase letters indicate significant differences between samples (Mann-Whitney U-test, *p* < 0.05; n ≥ 75 nuclei from at least 7 nodules). Colour code: purple: symbiotic NIN, green: non-symbiotic NIN, grey: NIN paralog.

Despite AtNLP2 being nitrate-dependent for nuclear localization, it was still able to partially function in symbiosis (Fig. 1). This could be due to the presence of a low amount of exogenous nitrate in the experiment (0.5 mM). To test whether the function of AtNLP2 in nodulation is nitrate-dependent, we examined the symbiotic functioning of AtNLP2 in absence of nitrate. *Mtnin-1* roots complemented with *AtNLP2* formed nodules under low nitrate (0.5 mM), but in absence of nitrate neither nodules nor infection threads were observed (Fig. 3c). This correlated with a lower amount of AtNLP2 in the nucleus under these conditions (Fig. 3d,e). In contrast, *Mtnin-1* roots transformed with *MtNIN* formed nodules in the absence of nitrate (Extended Data Fig. 6a). To assess whether enhanced nuclear localization could improve AtNLP2 function, we fused nuclear localization signals (NLS) to its termini and expressed these in *Mtnin-1*. Despite increased nuclear accumulation, these constructs failed to induce nodules (Extended Data Fig. 6b-d), likely due to interference of the NLS, as similar fusions to MtNIN also impaired its function (Extended Data Fig. 6e).

These data show that non-symbiotic NIN such as AtNLP2 requires nitrate to be nuclear localized and function in nodulation.

### A single amino acid adaptation makes AtNLP2 functionally similar to a symbiotic NIN

Previous studies showed that a specific phosphorylation site required for nitrate-triggered nuclear retention has been lost in legume NIN proteins^30,32–34^. Sequence analysis indicated that this is the result of lineage-specific events that occurred multiple times after NIN was recruited in nodulation, while it was maintained in multiple nodulating non-legume species such as Parasponia (Extended Data Fig. 7).

To identify changes enhancing symbiotic NIN function, we compared symbiotic NINs with non-symbiotic NIN orthologs across a wide range of plant species and identified seven residues specifically conserved in symbiotic NINs, suggesting positive selection in the NFC (Fig. 4a, Extended Data Fig. 7 and Supplementary Table 2). Among the identified motifs, position 1 and 5 have the potential to create an extra NLS for symbiotic NINs (Fig. 4a), whereas position 2, also identified by Zhang et al.^33^, is in the middle of the nitrate binding domain ^29^. Positions 5, 6, and 7 are relatively close to the DNA binding domain. Positions 4, 5, and 6 are within a region for which no specific structure is predicted (Fig. 4a,b).

**Fig. 4.**
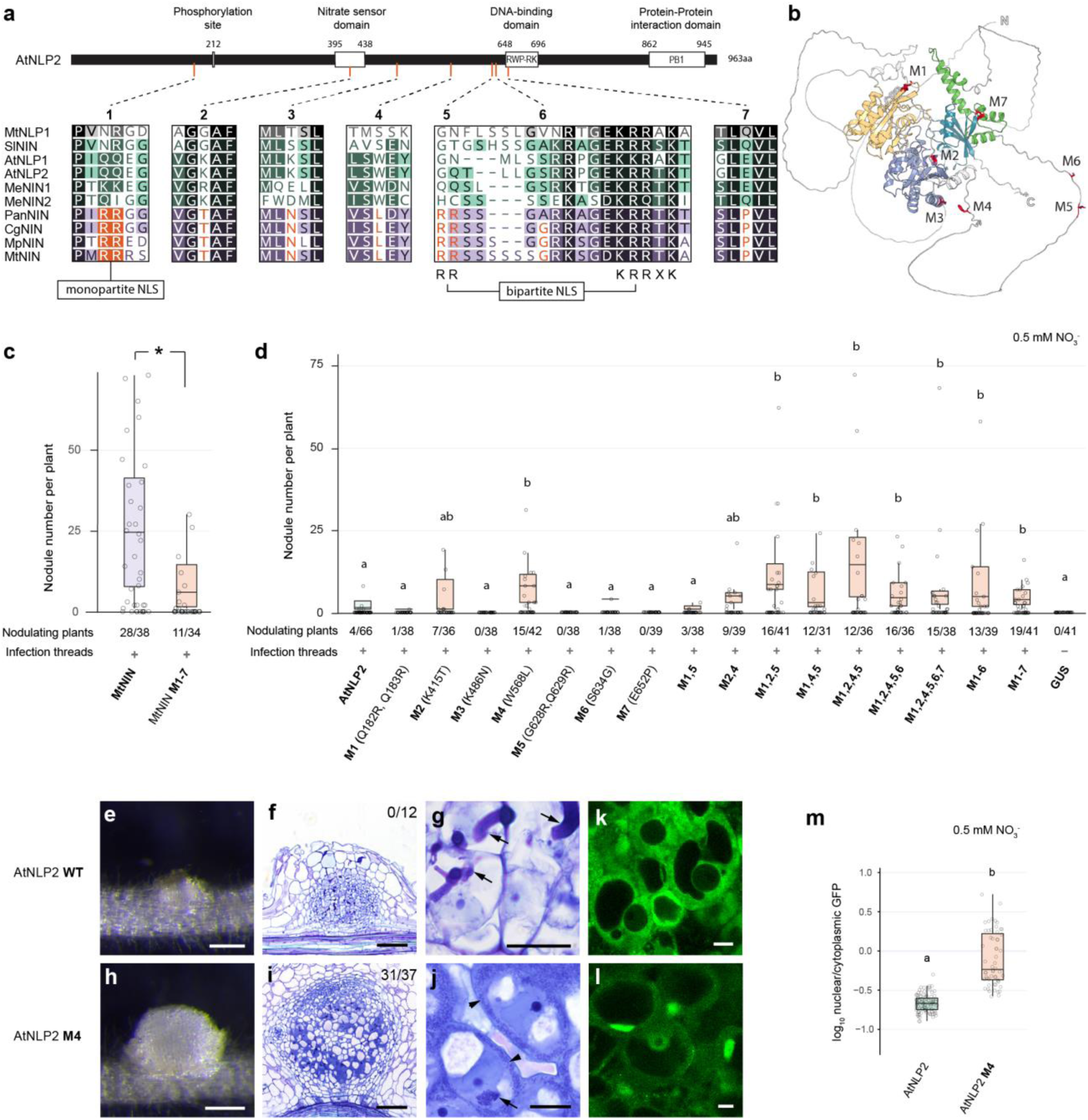
A single amino acid adaptation improves AtNLP2 functioning in nodulation. **a**, Schematic illustration of seven conserved amino acid changes between symbiotic (purple) and non-symbiotic (green) NIN orthologs. Mt: Medicago, Sl: tomato, At: Arabidopsis, Me: cassava, Pan: Parasponia, Cg: *Casuarina glauca*, Mp: *Mimosa pudica.* **b**, Structural prediction analysis of AtNLP2 forms three main domains; the N-terminal nitrate responsive domain that forms two globes (yellow and purple), an RWP-RK DNA-binding domain (blue) and a PB1 protein-protein interaction domain (green). The interdomain regions are highly disordered (white). The positions of the seven conserved amino acids (M1-M7) are marked in red. **c**, Number of nodules formed on *Mtnin-1* mutant roots complemented with *MtNIN* and *MtNIN^M^*^1–7^. Box plots show number of nodules per nodulated plant. Asterisk indicates significant difference (Mann-Whitney *U-*test, p < 0.05). **d**, Number of nodules formed on *Mtnin-1* mutant roots complemented with wildtype *AtNLP2* and *AtNLP2* with different adaptations at single or multiple positions. Lowercase letters indicate significant differences between samples (Kruskal-Wallis and post-hoc Dunn’s test, Benjamini-Yekutieli adjusted *p* < 0.05). **e**, **h**, Stereomicroscope images of nodules formed on *Mtnin-1* complemented with *AtNLP2* and *AtNLP2^M^*^4^. Scale bars: 2 mm. **f**, **i**, Longitudinal sections stained with toluidine blue. Numbers indicate the number of nodules with released bacteria per analyzed nodules. Scale bars: 100 μm. **g**, **j**, Magnification of nodule cells. Arrows indicate infection threads; arrowheads indicate released rhizobia. Scale bars: 20 μm. **k**, **l**, Confocal images showing the subcellular localization of wildtype AtNLP2 and AtNLP2^M4^ in Medicago nodule infection zone cells under 0.5 mM nitrate. Scale bars: 10 mm. **m**, Quantification of subcellular localization shown in (**k, l**). Lowercase letters indicate significant differences between samples (Kruskal-Wallis and post-hoc Dunn’s test, Benjamini-Yekutieli adjusted *p* < 0.05; n ≥ 50 nuclei from at least 7 nodules).

To test whether the identified amino acid changes represent important motifs for symbiotic NIN functioning, we introduced the corresponding AtNLP2 residues into MtNIN. This showed a significantly lower number of nodules formed by the mutated MtNIN^M1–7^ compared to MtNIN control, although both were able to form functional nodules (Fig. 4c, Extended Data Fig. 8). This implies that one or more of these seven residues are important for nodule formation.

To study whether single sites of amino acid changes can improve the function of non-symbiotic NIN, we introduced each of seven symbiotic-NIN-specific residues into AtNLP2 and assessed their function in trans-complementation of *Mtnin-1*. Compared to wildtype AtNLP2, AtNLP2^M^^4^ (W568L) significantly increased its complementation efficiency, while the others did not (Fig. 4d). AtNLP2^M4^ not only induced the formation of more nodules, these nodules were also markedly bigger. Sections of these nodules showed bacteria were released into nodule cells. Further, a meristem was formed at the apex, and vasculature in peripheral tissue. This nodule phenotype is similar to that of the nodules formed on *Mtnin-1* roots trans-complemented by *PanNIN* (Fig. 4e-j, Extended Data Fig. 9).

To investigate whether the enhanced functioning of AtNLP2^M4^ in nodulation is linked to increased nuclear localization, we compared the subcellular localization of *GFP-AtNLP2^M^*^4^ and *GFP-AtNLP2* in Medicago nodules. *GFP-AtNLP2^M^*^4^ showed increased nuclear localization when compared to *GFP-AtNLP2* (Fig. 4k-m), suggesting that this modification likely enhances AtNLP2 function by promoting its nuclear localization.

Next, we tested whether introducing multiple mutations into AtNLP2 would further improve its function in nodule formation. Some combinations significantly improved complementation efficiency over wild-type AtNLP2, but not beyond AtNLP2^M^^4^ (Fig. 4d). Interestingly, AtNLP2^M^^1,2,5^, without mutating position 4, also significantly enhanced nodule formation, although no rhizobial release was observed (Extended Data Fig. 10). These data demonstrate that multiple ways of amino acid modifications can improve NIN functioning in nodule symbiosis, providing alternative evolutionary trajectories enhancing symbiotic NIN efficiency.

We also tested whether the identified changes improved nodulation at zero nitrate growth conditions. This showed that none of the AtNLP2 mutant variants, including AtNLP2^M4^, complemented *Mtnin-1* for nodulation (Extended Data Fig. 11). This demonstrates that a low concentration of exogenous nitrate is essential for functioning of AtNLP2, AtNLP2^M4^, and other AtNLP2 mutant variants, whereas this is not the case for symbiotic NINs.

Given the efficient functioning of AtNLP2^M4^ in nodulation, we further investigated the evolutionary trajectory of position 4. It shows that the leucine residue at this position likely did not arise at the base of the NFC but instead represents the most probable ancestral state (Extended Data Fig. 7). Non-nodulating plant species such as *Petunia axillaris* and *Vitis vinifera* both possess a NIN ortholog with a leucine at position 4. However, these proteins did not trigger nodule primordium formation when introduced into Mt*nin-1*, and only some infection threads were formed in case of VvNIN (Extended Data Fig. 12). This shows that a leucine at position 4 alone is not sufficient to induce nodule formation. Comparison of the region surrounding position 4 of NINs with paralogues showed that it is unique to the NIN subclade. Prior to the evolution of the NFC, both position 4 and its surrounding region were variable (Extended Data Fig. 7). In contrast, they became highly conserved in symbiotic NINs within NFC, highlighting strong evolutionary pressure to maintain this region and in line with its functional significance in nodulation.

## Discussion

In this study, we uncovered an intriguing aspect of the evolution of symbiotic NIN. Although its recruitment into symbiosis dramatically shifted its function from a nitrate sensor to a transcriptional response factor in nodulation, non-symbiotic NINs were already partially capable of performing this function. Non-symbiotic NIN orthologs from a variety of species can induce symbiotic gene expression, enabling infection thread formation, and the initiation of nodule organogenesis. However, they do this with low efficiency and without the ability to facilitate the intracellular colonization of nodule cells by rhizobia. Remarkably, a single amino acid change is sufficient to convert a non-symbiotic NIN, AtNLP2, into a NIN protein functionally similar to the symbiotic PanNIN, including the ability to facilitate bacterial intracellular accommodation in nodules.

The single amino acid change from tryptophane to leucine at position 4 enhances the nuclear localization of AtNLP2. Possibly by causing a conformational shift that either exposes the nuclear localization signal or masks the nuclear export signal. However, for AtNLP2^M4^ to function in nodulation, some nitrate must be present, indicating that a nitrate-dependent activation is still required. In AtNLP7, nitrate triggers a conformational change that derepresses the protein via its N-terminus^29^. Due to current limitations in protein modelling an experimental determined structure of NIN will be essential to provide insights into the nitrate dependent activity.

The properties of non-symbiotic NINs in a nodulation context are variable with respect to their subcellular localization in response to nitrate, their ability to induce symbiotic gene expression, and their ability to induce rhizobium infection and nodule formation. This variability may result from relaxed selection pressure following the duplication of an NLP within eudicots, giving rise to the NIN and NLP1 clades^16^. Lineage-specific duplication may allow further functional diversification as observed in Arabidopsis and cassava NIN orthologs. In our study, AtNLP2 was the non-symbiotic NIN with the best ability to function in nodule formation. The observation that AtNLP2 can form nodule primordia demonstrates that this symbiotic functionality can evolve without nodulation as selection pressure. A single amino acid change that restores the most probable ancestral state of position 4 further improves the symbiotic functionality of AtNLP2. Furthermore, we used ancestral sequence reconstruction to filter out lineage-specific evolution, though potential mispredictions may underestimate true ancestral functionality. The primordia induced by the resurrected NIN_NFC_ support that the common ancestor of the NFC possessed a NIN protein able to function in nodulation. Taken together, these findings support the hypothesis that NIN was predisposed to function in nodulation at the time of its recruitment into nitrogen-fixing nodule symbiosis.

Our study revealed the ability of non-symbiotic NINs to function in nodulation. Furthermore, we identified amino acids in NIN that are under strong selection pressure in the NFC and are important for its functionality in symbiosis after its recruitment. These findings provide critical insights into the molecular mechanisms driving the evolution of nodulation and could inform strategies to engineer nodulation in non-legume crops.

## Methods

### Plant material and growth conditions

The *Medicago truncatula* (Medicago) *nin-1* knock-out mutant (in Jemalong A17 background), and wildtype Jemalong A17 plants were used in this study. *Agrobacterium rhizogenes* strain MSU440 was used for transformation, as described previously^35^. The composite plants were grown at 21°C and 16h light/8h dark regime, in perlite saturated with Färhaeus medium, with Ca (NO_3_)_2_ added to the indicated NO_3_^-^ concentrations. For nodulation experiments, plants were first grown in perlite for one week, then inoculated with *Sinorhizobium meliloti 2011* rhizobia constitutively expressing GFP (OD_600_=0.1, 2 mL per plant), and harvested four weeks post inoculation. Transgenic roots were selected for analysis based on expression of the visual selection marker mCherry. For the trans-activation assay, plants were grown for 3 weeks on perlite saturated with Färhaeus medium, without NO_3_^-^ added. Then, plants were treated with Färhaeus containing 10 µM dexamethasone (from a 10 mM stock in DMSO) or containing 0.1% DMSO as a mock treatment. The transgenic roots were harvested 16 hours after treatment. For the subcellular localization studies, plants were grown in perlite saturated with Färhaeus medium, without NO_3_^-^. After growing in perlite for one week, the plants were inoculated with wildtype *S. meliloti 2011* rhizobia (OD_600_=0.1, 2 mL per plant). After four weeks post inoculation, the nodules were harvested for further analysis.

### Constructs

The constructs used in this study were assembled by Golden Gate cloning. Standard promoters, terminators, backbones, and binary vectors were obtained from the MoClo Toolkit and MoClo Plant Parts Kit ^36,37^, Addgene Kit #1000000044, and Kit #1000000047). Modules specific for this study were de novo synthesized. Supplementary Table 4 lists the sequences of all used Golden Gate cloning vectors.

To analyse the subcellular localization of different NINs, non-symbiotic NINs and MtNLP1 fused to GFP were driven by the constitutive *Lotus japonicus UBIQUITIN1* promoter (*pLjUBQ1*) ^38^. When using this promoter to drive the expression of symbiotic *MtNIN* and *PanNIN* N-terminal *GFP* fusions, nodule formation was suppressed, likely due to autoregulation of nodulation controlled by symbiotic NIN ^6^. Therefore, we used the nodule-specific *MtNIN* promoter to drive GFP-tagged *MtNIN* and *PanNIN*. Related constructs are included in Supplementary Table 4.

### Microscopy

The images of the nodules and the roots were taken using a stereomicroscope (M165 FC, Leica). For the section images, embedding of plant tissue in plastic, sectioning and tissue staining were performed as described previously^39^. Sections were analyzed using a DM5500B microscope equipped with a DFC425C camera (Leica). The subcellular localization and infection threads were studied using a Leica SP8 confocal microscope.

To analyse the subcellular localization under different nitrate concentrations, nodules were hand-sectioned and treated with 0 mM, 0.5 mM or 20 mM KNO_3_ solution for 1 hour before microscopy. The GFP intensity in the nucleus and cytoplasm was quantified using ImageJ software (1:53c) on the confocal pictures.

### RNA isolation, cDNA synthesis, and qRT-PCR

RNA was isolated from Medicago roots using the EZNA Plant RNA kit (Omega Bio-Tek), following manufacturer’s protocol. The iScript cDNA synthesis kit (Bio-Rad) was used to synthesize cDNA with a blend of oligo (dT) and random primers. Primers targeting symbiotic marker genes are listed in Supplementary Table 3. qRT-PCR was performed on the CFX96 connect Real-Time PCR system (Bio-Rad), using SYBR Green Supermix (Bio-Rad). The gene expression was normalized using *MtACTIN2* as a reference gene.

### Electro mobility shift assay (EMSA)

Proteins for EMSA were produced using the TNT SP6 High-Yield Wheat Germ Expression System (Promega) using the vectors pL1M-R1 EMSA 6HA-MBP-MtNINct, 6HA-MBP-PanNINct, 6HA-MBP-MesNIN1ct, 6HA-MBP-MesNIN2ct, 6HA-MBP-SlNINct, 6HA-MBP-AtNLP2ct, 6HA-MBP-AtNLP1ct, 6HA-MBP-MtNLP1ct (Supplementary Table 4, Extended Data Fig. 3c) as input. The in vitro translation was incubated for 2.5 hours at 25 °C. Fluorescent probes were generated by PCR on probe template vectors pL0M-P NINBS MtCEP7, pL0M-P NINBS MtCLE13, pL0M-P NINBS MtLBD16, pL0M-P NINBS MtNF-YA1 or pL0M-P NINBS MtNF-YB16 (Supplementary Table 4) using primers conjugated to fluorescent dye IR700 (Supplementary Table 3). PCR fragments were purified using the GeneJet PCR purification kit (Thermo Scientific). 4 µl of the in vitro translation mix and 20ng of the PCR probe were combined in EMSA binding buffer to a total volume of 12 µl (0,25 mg/ml BSA, 7,5 mM HEPES-NaOH pH 7.3, 0,7 µM DTT, 70 mM KCl, 1,5mM MgCl_2_, 60 µg/ml salmon sperm DNA, 2,5% CHAPS, 8% glycerol), and incubated for 30 minutes on ice. The reaction was run on a 0,5x TBE, 5% acryl-bisacrylamide gel, and visualized using a LiCor Odyssey fluorescence gel-scanner. The intensity of bands was analyzed using ImageJ software (1:53c).

### Western blot analysis

To visualize the synthesized protein used in EMSA, Mini-PROTEAN TGX Stain-Free Gels (Bio-Rad) were used for running the protein gels, then the samples were transferred to polyvinylidene difluoride membranes. After blocking with 3% bovine serum albumin, 5000 times diluted anti-HA-HRP (Miltenyi Biotec) with the ECL Western Blotting Substrate (Bio-Rad) were used for detection.

### Alignment and comparison of amino acid sequences, ancestral sequence reconstruction

Protein sequences of NIN and NLP1 orthologs were identified by BLAST, and their orthology to MtNIN and MtNLP1 was confirmed based on their clustering with these proteins in a phylogenetic tree (Extended Data Fig. 7). Protein sequences and accession numbers are listed in Supplementary Table 5. A multiple sequence alignment was performed using the MAFFT alignment tool (1.2.2) in Geneious Prime software (GraphPad Software LLC). A phylogenetic tree was built using MrBayes (3.2.7a x86_64), using AtNLP7 as an outgroup, and without topology restraints.

To quantify conservation of amino acids in subgroups, the total alignment was subsetted to contain only the NFC or all dicot NIN orthologs. Jalview (2.11.4.1; The Barton Group, University of Dundee) was used to evaluate conservation levels of the subsets (Supplementary Table 2). The seven identified sites were selected based on their high conservation in NFC but low conservation across all dicot NIN orthologs, combined with manual evaluation.

For ancestral sequence reconstruction, we used the tree built using MrBayes without topology constraints as a starting point to select the nodes for NIN reconstruction; the NFC clade (NIN_NFC_), and the most recent non-symbiotic ancestor, encompassing the NFC as well as species belonging to both Fabids and Malvids (NIN_Rosids_). We used this tree and the alignment as input for GRASP (version 21-Mar-2024), using two independent methods of indel prediction; bi-directional edge parsimony (BEP) and simple indel-coding maximum likelihood (SICML). In parallel, we used MrBayes to reconstruct the ancestors, setting the nodes to be reconstructed as constraints. We compared the ancestors obtained by the three different methods and manually inspected and curated each position where the models disagree (Supplementary Table 2).

### Transient expression in *Nicotiana benthamiana* leaves

*Nicotiana benthamiana* (tobacco) were grown in soil for 4 weeks. The plants were treated with 20 mM KNO_3_ or water (mock) one day before the infiltration. *Agrobacterium tumefaciens* strain C58 carrying constructs were co-infiltrated with silencing inhibitor P19 in tobacco leaves as described previously^40^. The infiltrated leaves were collected for confocal microscopy three days post infiltration.

### Statistical analysis

All statistical analyses were performed using R-studio software (Posit Software, PBC). Nodule number per plant, gene expression in transactivation assay, and ratio of fluorescence in the nucleus and cytoplasm were not normally distributed (Shapiro-Wilk p < 0.05). Therefore, non-parametric tests were used in all analyses. For multiple comparisons, we used the Kruskall-Wallis test, with Dunn’s test post hoc analysis. *p*-values were corrected for multiple testing using the Benjamini-Yekutieli method. For single or pairwise comparisons, the Mann-Withney *U*-test was used. Bound fraction in EMSA assays was normally distributed (Shapiro-Wilk p > 0.05), and were analzed using a Student’s t-test versus control samples, with multiple testing correction using the Benjamini-Yekutieli method.

### NIN structure modeling

All models were predicted using Alphafold 3 with settings unaltered from the default, and with all shown models being representative of five simulations.

## Supporting information

Comparison of codon usage between different NIN orthologs and the Medicago truncatula genome

Ancestral sequence reconstruction and analysis of amino acid conservation

Constructs used in this study

Protein sequences used for analysis

## Data availability

All data are available within this article and its Supplementary Information.

## Acknowledgments

We thank Tom Peeters, Cloé Villard, Robin van Velzen, Joël Klein, Joan Wellink, Ella Köbben, and Stan van Wijk for their contribution and suggestions to this project. This project is supported by funding from the project Enabling Nutrient Symbioses in Agriculture (ENSA) that is funded by Bill & Melinda Gates Agricultural Innovations (INV-57461), China Scholarship Council (201506300062 to J.L., 201906170085 to S.Y., 202008150090 to M.L.), and the Dutch Science Organization (Nederlandse Organisatie voor Wetenschappelijk Onderzoek VI.Veni.212.132) to R.H.

## Author contributions

J.L. conceived and designed the work, performed the experiments, analysed the data, and wrote the manuscript. S.Y. conceived and designed the work, performed the experiments, analysed the data, and wrote the manuscript. M.L. performed experiments and analysed the data. D.S. performed experiments. M.T. performed experiments. R.B. performed experiments and analysed the data. K.R.A analysed the data. F.V. performed experiments. O.K. analysed the data. R.G. conceived and designed the work, analysed the data and wrote the manuscript. T.B. conceived and designed the work, analysed the data and wrote the manuscript. R.H. conceived and designed the work, performed the experiments, analysed the data, and wrote the manuscript.

## Competing interests

Some findings in this manuscript are considered for patent application.

---

**Extended Data Fig. 1.**
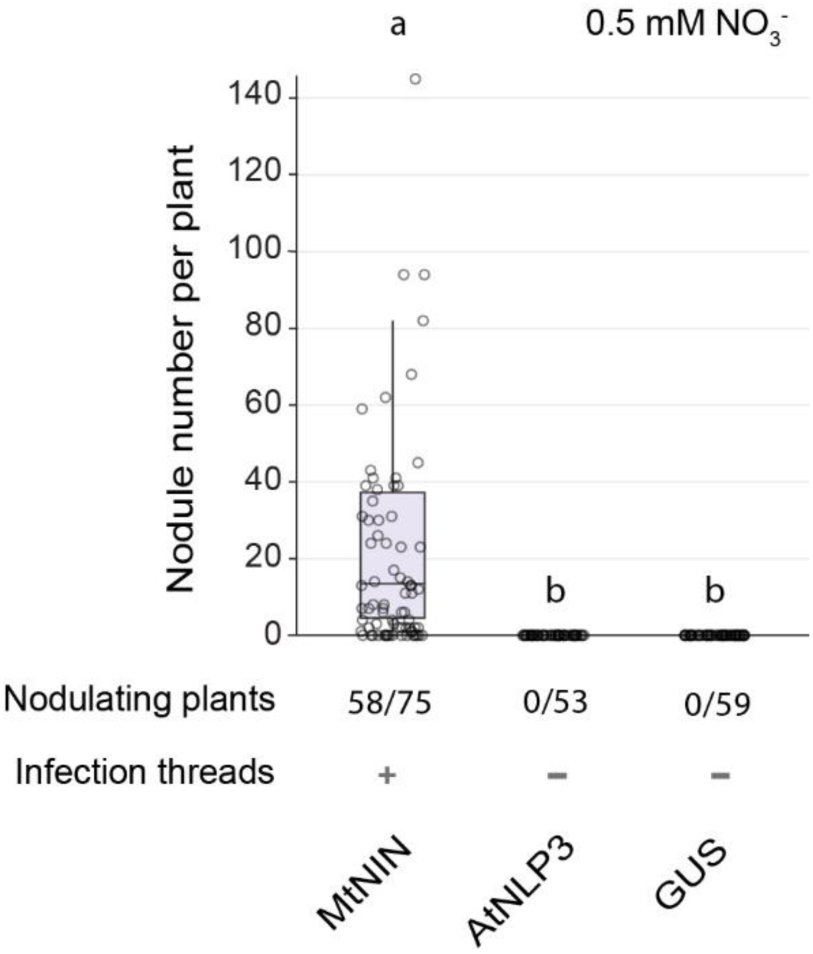
AtNLP3 is not functional in root nodule symbiosis. Number of nodules formed on *Mtnin-1* mutant roots complemented with MtNIN and AtNLP3. Plants were harvested at 4 weeks post inoculation with *S. meliloti* 2011 expressing GFP. Box plots show the number of nodules per nodulated plant. Lowercase letters indicate significant differences between samples (Kruskal-Wallis and post-hoc Dunn’s test, Benjamini-Yekutieli adjusted p < 0.05).

**Extended Data Fig. 2.**
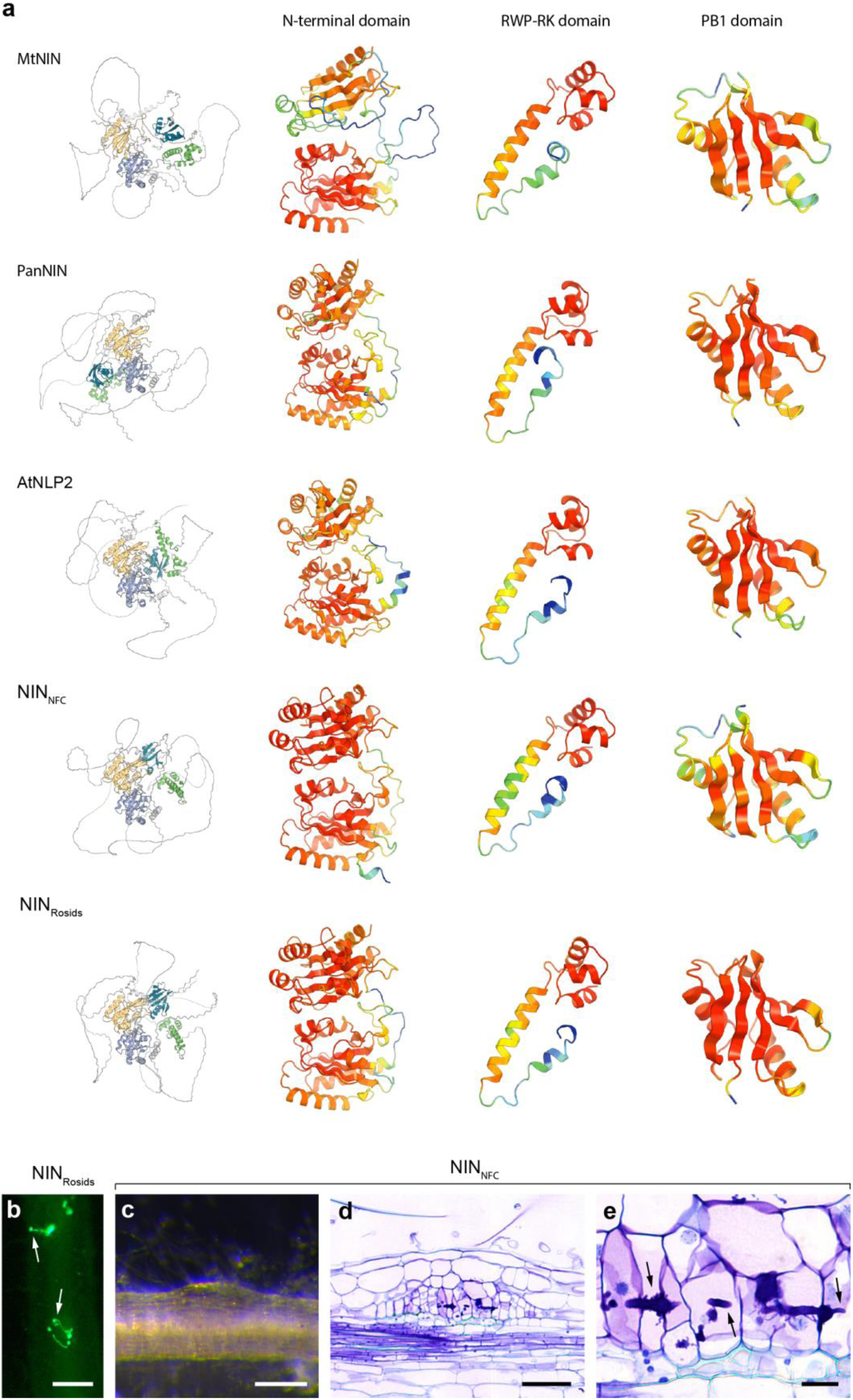
Functionality of resurrected ancestral NIN proteins. **a**, Predicted models of extant NINs and resurrected ancestors share structural conserved domains. Overview of full-length NIN and three conserved domains; N-terminal domain, RWP-RK DNA-binding domain, and PB1 protein-protein interaction domain, colored by their pLDDT score, with red indicating a score of 90 and higher. **b**, Green fluorescence stereomicroscopy images showing infection threads (arrows) formed on *Mtnin-1* roots complemented with the resurrected NIN_Rosids_ ancestor. Scale bars: 2 mm. **c**, Stereomicroscope images showing a nodule primordium formed on *Mtnin-1* roots transformed with the NIN_NFC_ ancestor. Scale bar: 2 mm. **d**, Longitudinal section of NIN_NFC_ induced nodule primordium, stained with toluidine blue. Scale bar: 100 μm. **e**, Magnification of (**d**). Arrows indicate infection threads. Scale bar: 20 μm.

**Extended Data Fig. 3.**
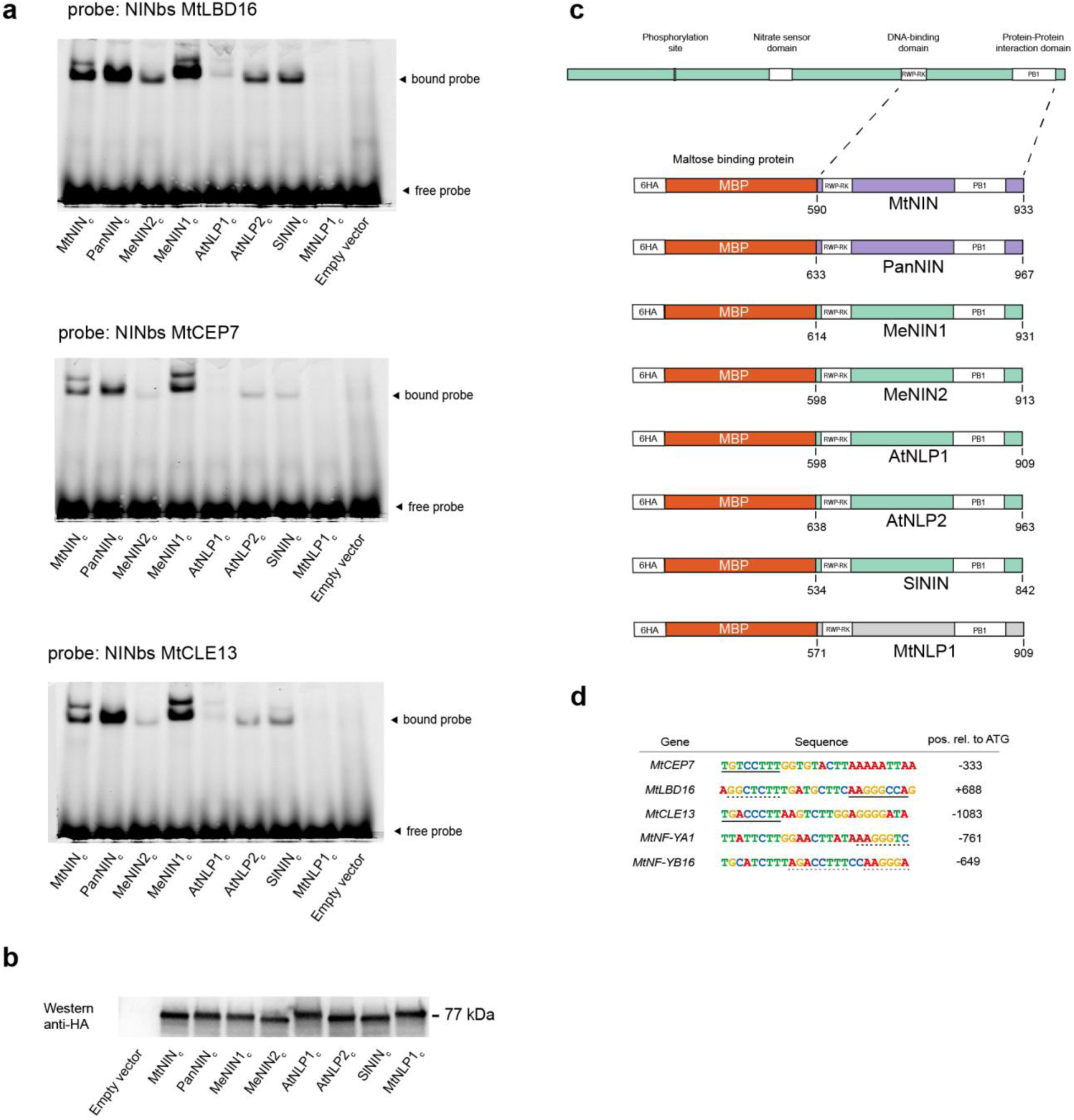
C-terminus of different NIN-orthologs can bind to symbiotic NIN-binding sites *in vitro*. **a**, Electrophoretic mobility shift assay (EMSA) testing the binding of the C-terminus of different NIN-orthologs, fused with 6xHAtag-MBP, to fluorescently labelled NIN *cis*-regulatory binding sites (NINbs) of *MtLBD16, MtCEP7,* and *MtCLE13*. **b**, Western blot showing the relative concentration of different fusion proteins used in EMSA. **c**, Constructs used in the EMSA (Fig. 2a, Extended Data Fig. 3a). The C-terminal region of different NIN orthologs was used, which includes the DNA-binding domain (RWP-RK) and a protein-protein interaction domain (PB1). Numbers refer to the amino acid range of each NIN protein that was used. The NIN C-terminal region was fused to the maltose binding protein (MBP) to aid in solubility, and 6 repetitions of the hemagglutinin tag (6HA) for detection. **d**, Probe sequences that were used in EMSA (Fig. 2a, Extended Data Fig. 3a). Solid lines indicate full match to the AtNLP2 binding site described by ^18^, dashed lines indicate partial match. The position relative to the ATG start codon is indicated. Note that the NIN binding site of MtLBD16 is in the first intron ^26^.

**Extended Data Fig. 4.**
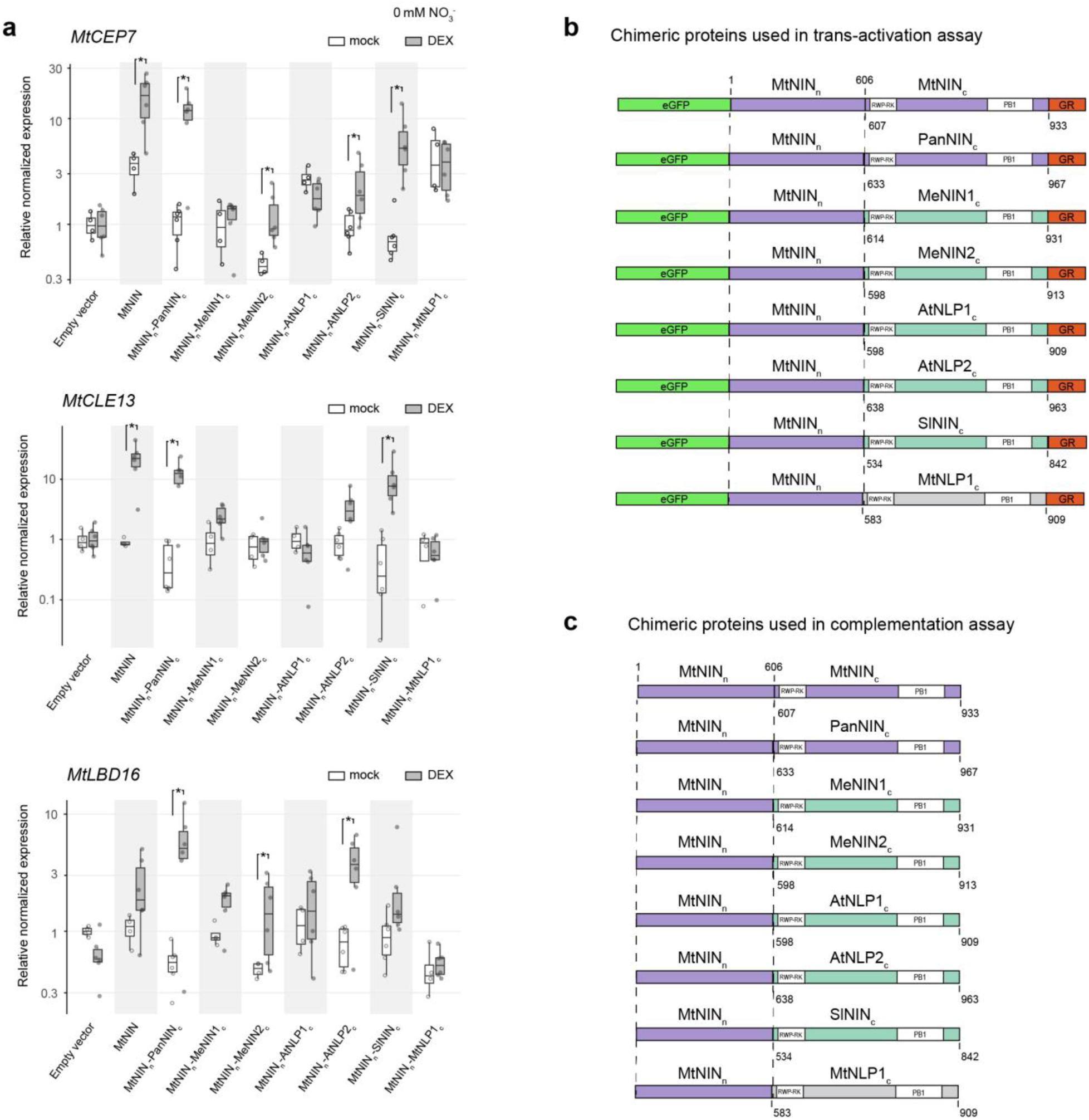
Symbiotic functionality of chimeric NIN proteins. **a**, qRT-PCR showing chimeric NIN proteins induce *MtCEP7*, *MtCLE13* and *MtLBD16* expression in a transactivation assay. Medicago roots producing the N-terminus of MtNIN fused to the C-terminus of different NIN/NLPs and the rat glucocorticoid receptor (Extended Data Fig. 4b), were treated with 10 µM dexamethasone (DEX) or DMSO (mock) for 16 hours. Expression levels were normalized to the average expression of mock treated empty vector roots. Asterisks indicate significant differences (Mann-Whitney U-test, p <0.05). **b**, Chimeric NIN proteins used in the transactivation assay (Fig. 2b, Extended Data Fig. 4a). The N-terminal region (residue 1-606) of MtNIN was fused to the C-terminal region of different NIN orthologs, which includes the RWP-RK domain and the PB1 domain. Numbers below each C-terminus show the amino acid range of each protein that was used. GR indicates the rat glucocorticoid receptor. **c**, Chimeric NIN proteins used in the complementation assay (Fig. 2c,d). The N-terminal region (residues 1-606) of MtNIN was fused to the C-terminal region of different NIN-orthologs, which includes the RWP-RK domain and the PB1 domain. Numbers below each C-terminus show the amino acid range of each protein that was used.

**Extended Data Fig. 5.**
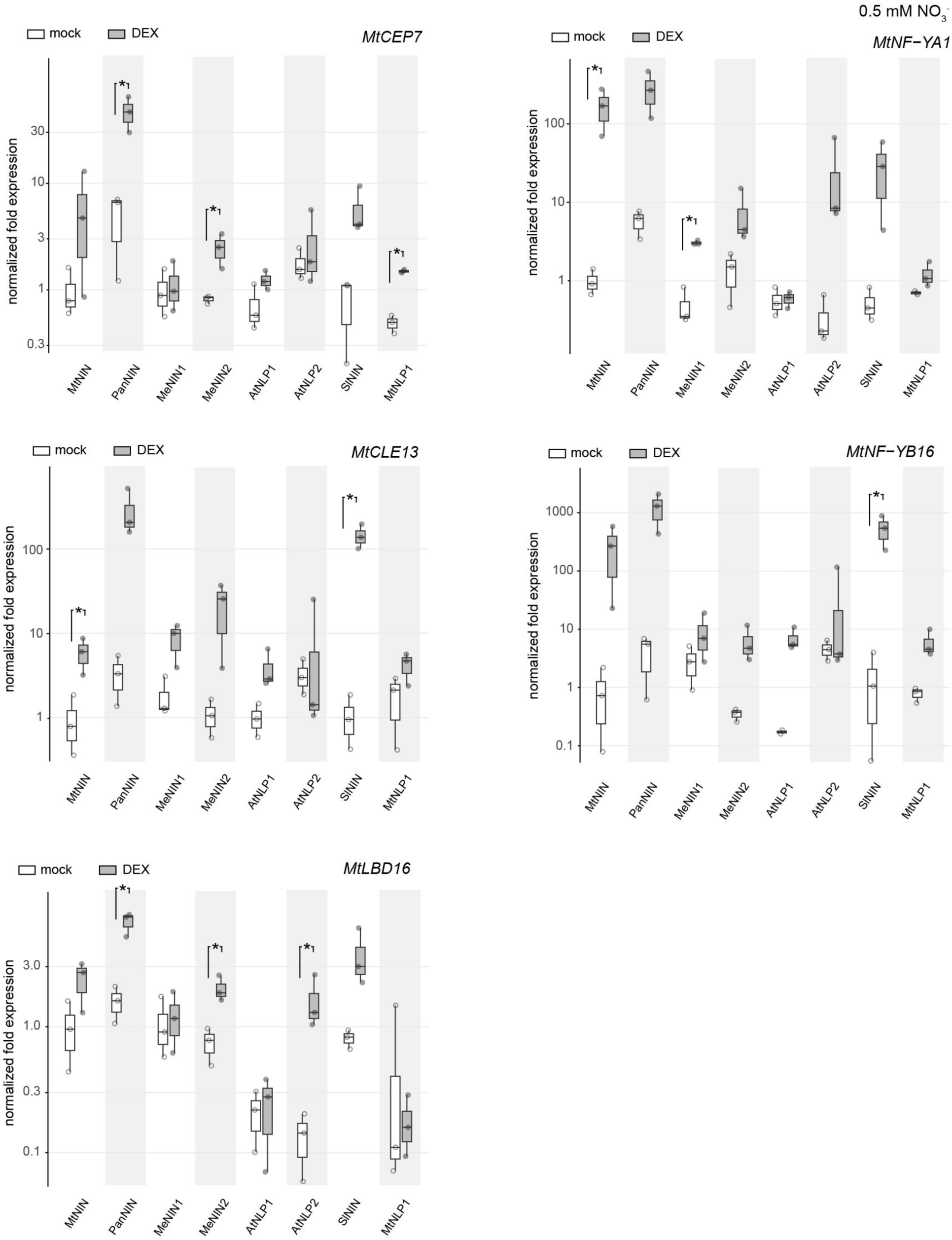
Transactivation assay using non-chimeric NIN-GR fusions. qRT-PCR showing induction of NIN target genes by non-chimeric NIN proteins fused to GR. Medicago roots expressing the constructs, were treated with 10 µM dexamethasone (DEX) or DMSO (mock) for 16 hours. Expression levels were normalized to the average expression of mock-treated roots expressing MtNIN-GR. Asterisks indicate significant differences (Student’s t-test, p < 0.05).

**Extended Data Fig. 6.**
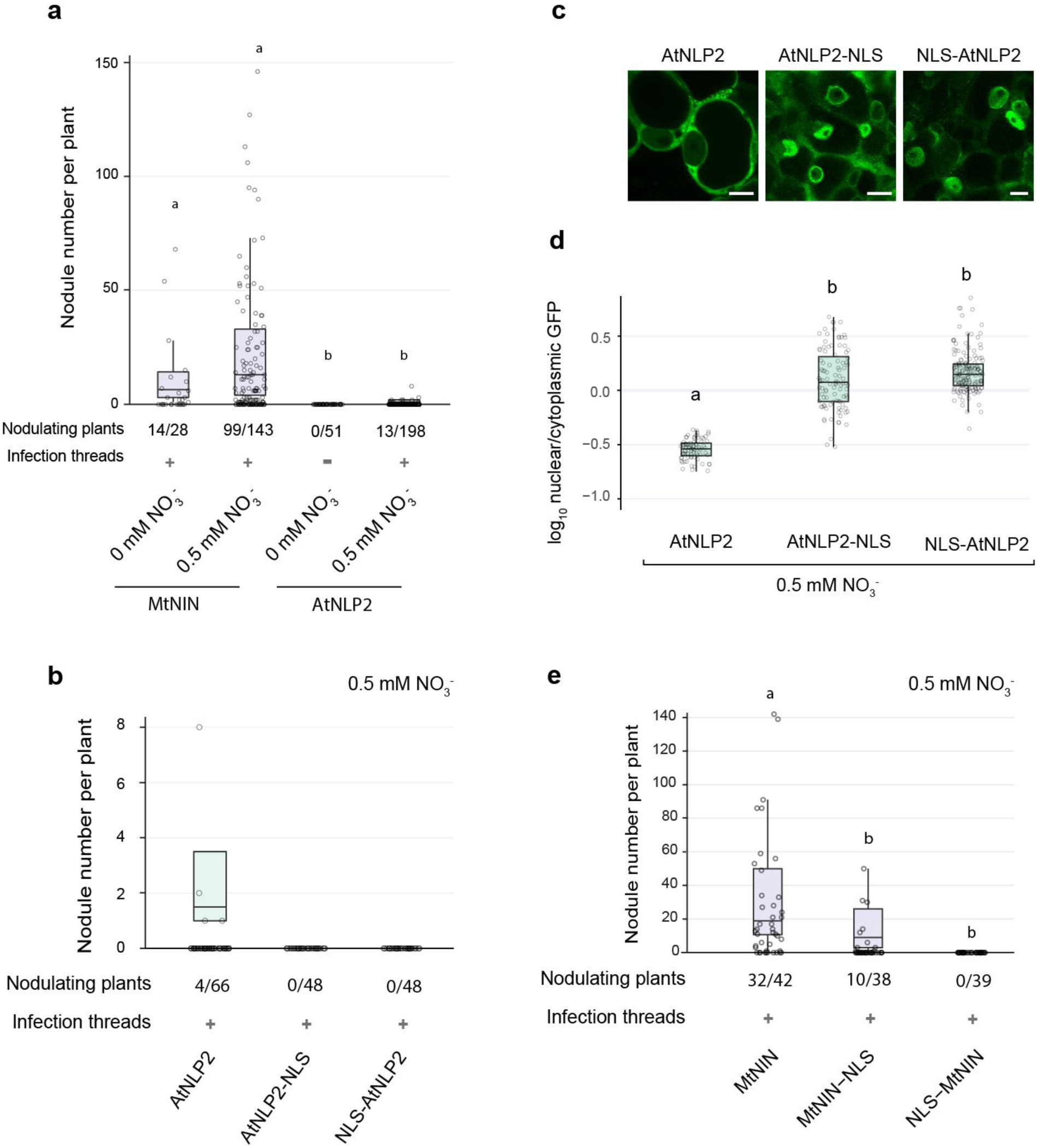
MtNIN, but not AtNLP2, can function in nodulation in absence of nitrate. **a**, Number of nodules formed on *Mtnin-1* mutant roots complemented with MtNIN or AtNLP2 under different exogenous nitrate concentrations. Plants were harvested at 4 weeks post inoculation with *S. meliloti* 2011 expressing GFP. Box plots show the number of nodules per nodulated plant. Lowercase letters indicate significant differences between samples (Kruskal-Wallis and post-hoc Dunn’s test, Benjamini-Yekutieli adjusted *p* < 0.05). **b**, Number of nodules formed on *Mtnin-1* mutant roots complemented with *AtNLP2* or *AtNLP2* fused with nuclear localization signals (NLS) at 0.5 mM nitrate. Differences are not significant (Kruskal-Wallis and post-hoc Dunn’s test, Benjamini-Yekutieli adjusted p < 0.05). **c**, Confocal images showing the subcellular localization of GFP-tagged AtNLP2 or AtNLP2 fused with nuclear localization signals (NLS) at 0.5 mM nitrate. Scale bars: 10 µm. **d**, Quantification of subcellular localization in (**c**). Lowercase letters indicate significant differences between samples (Kruskal-Wallis and post-hoc Dunn’s test, Benjamini-Yekutieli adjusted, p < 0.05; n ≥ 75 nuclei from at least 7 nodules). **e**, Number of nodules formed on *Mtnin-1* mutant roots complemented with MtNIN or MtNIN fused with nuclear localization signals (NLS) at 0.5 mM nitrate. Lowercase letters indicate significant differences between samples (Kruskal-Wallis and post-hoc Dunn’s test, Benjamini-Yekutieli adjusted p < 0.05).

**Extended Data Fig. 7.**
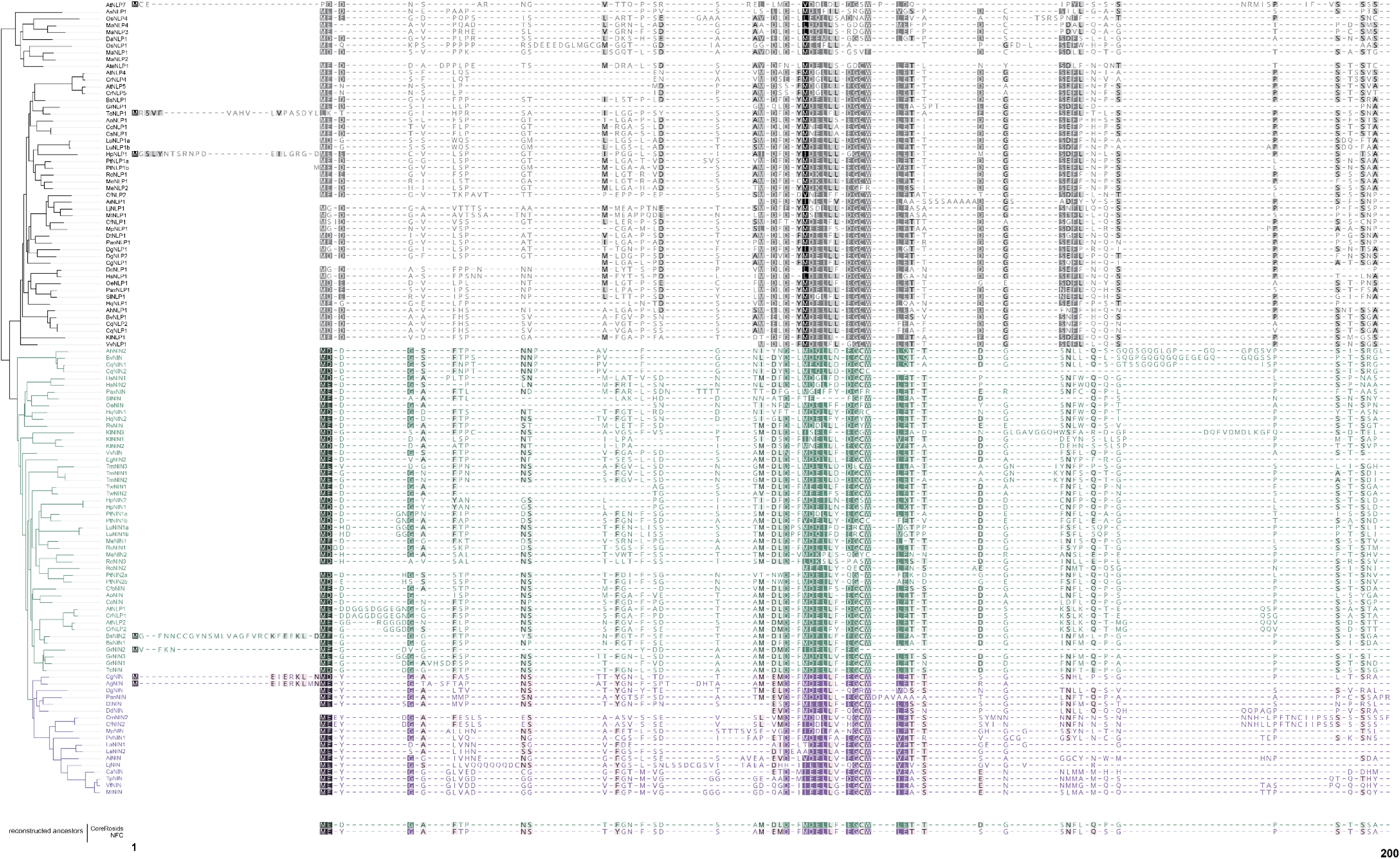
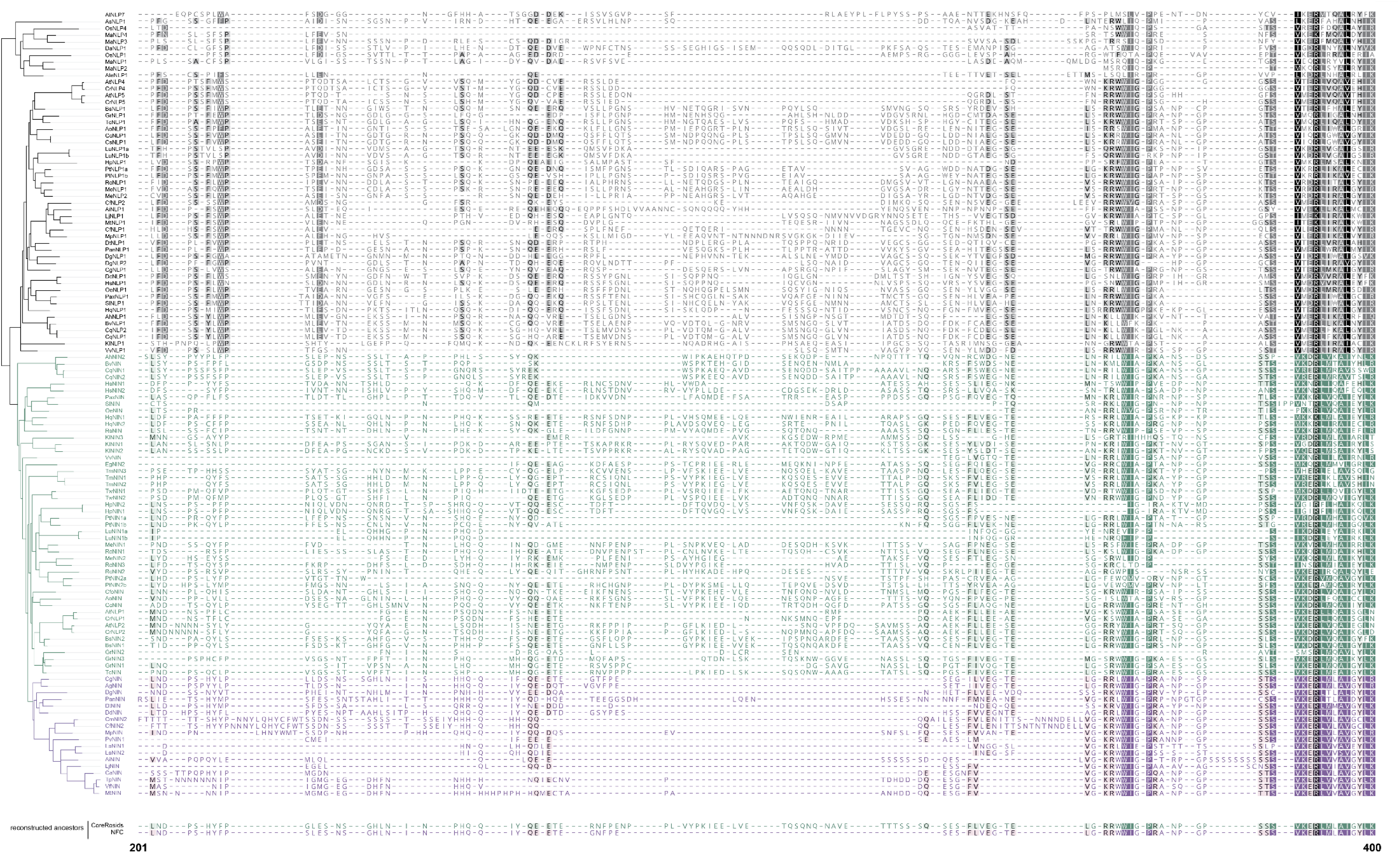
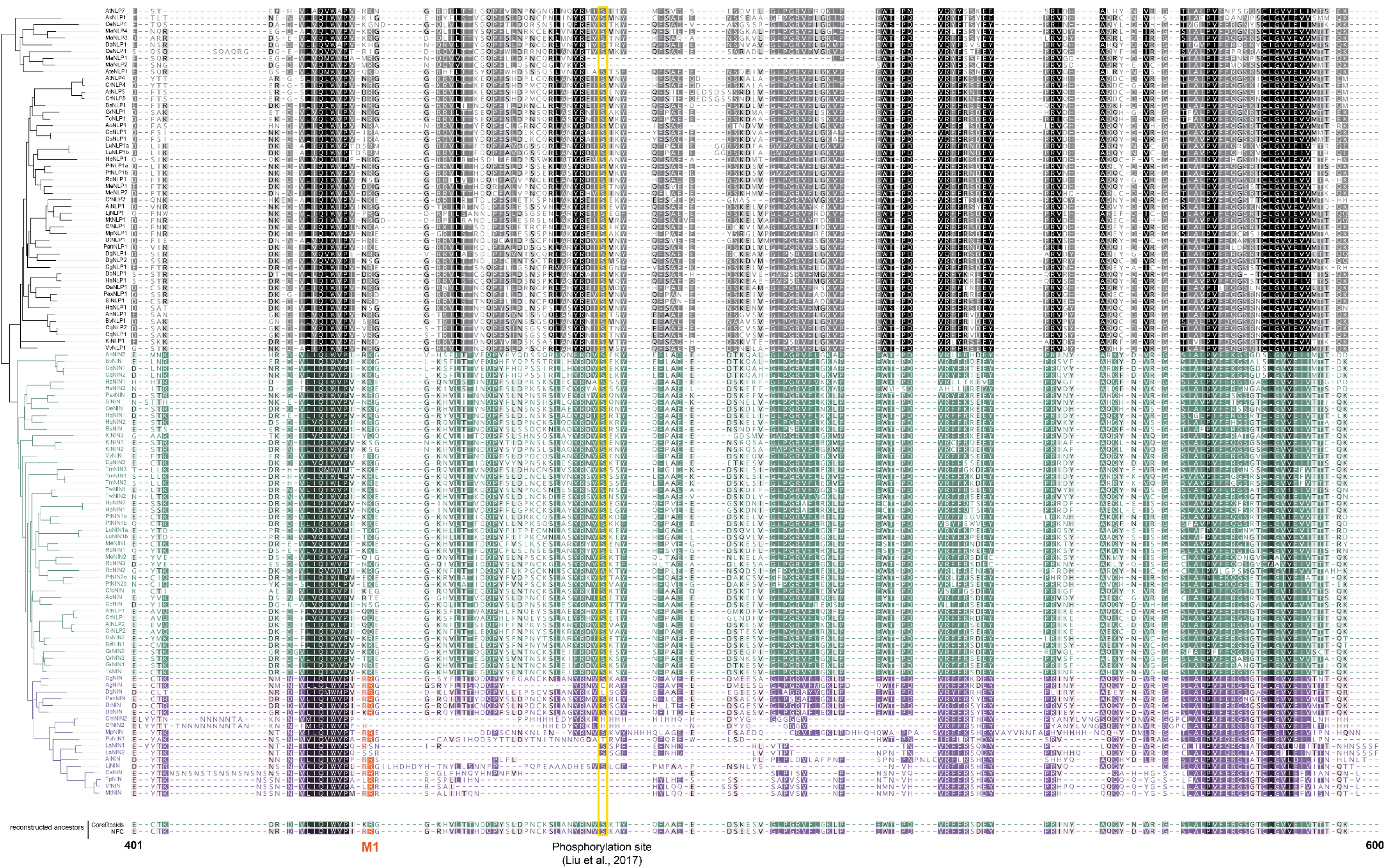
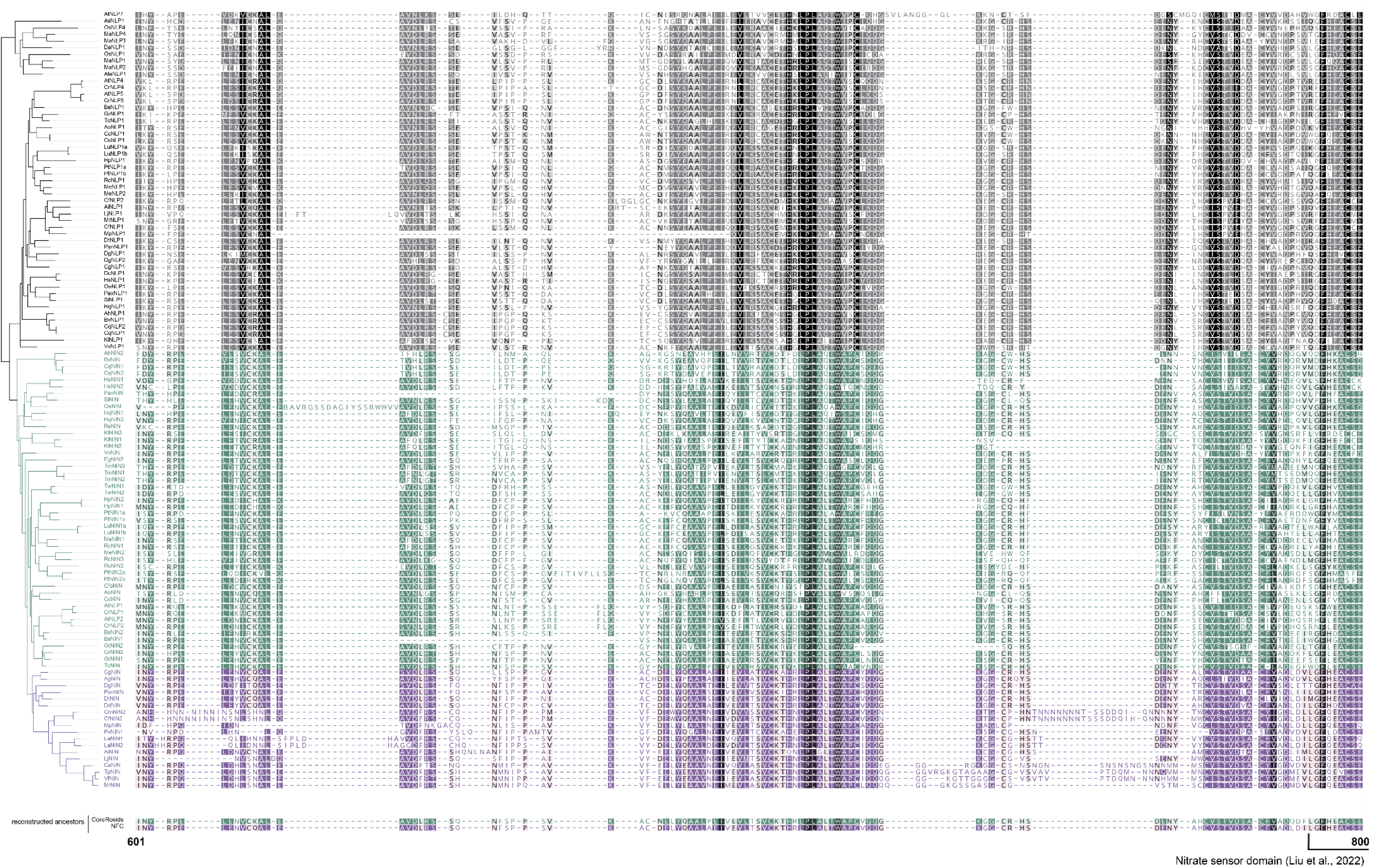
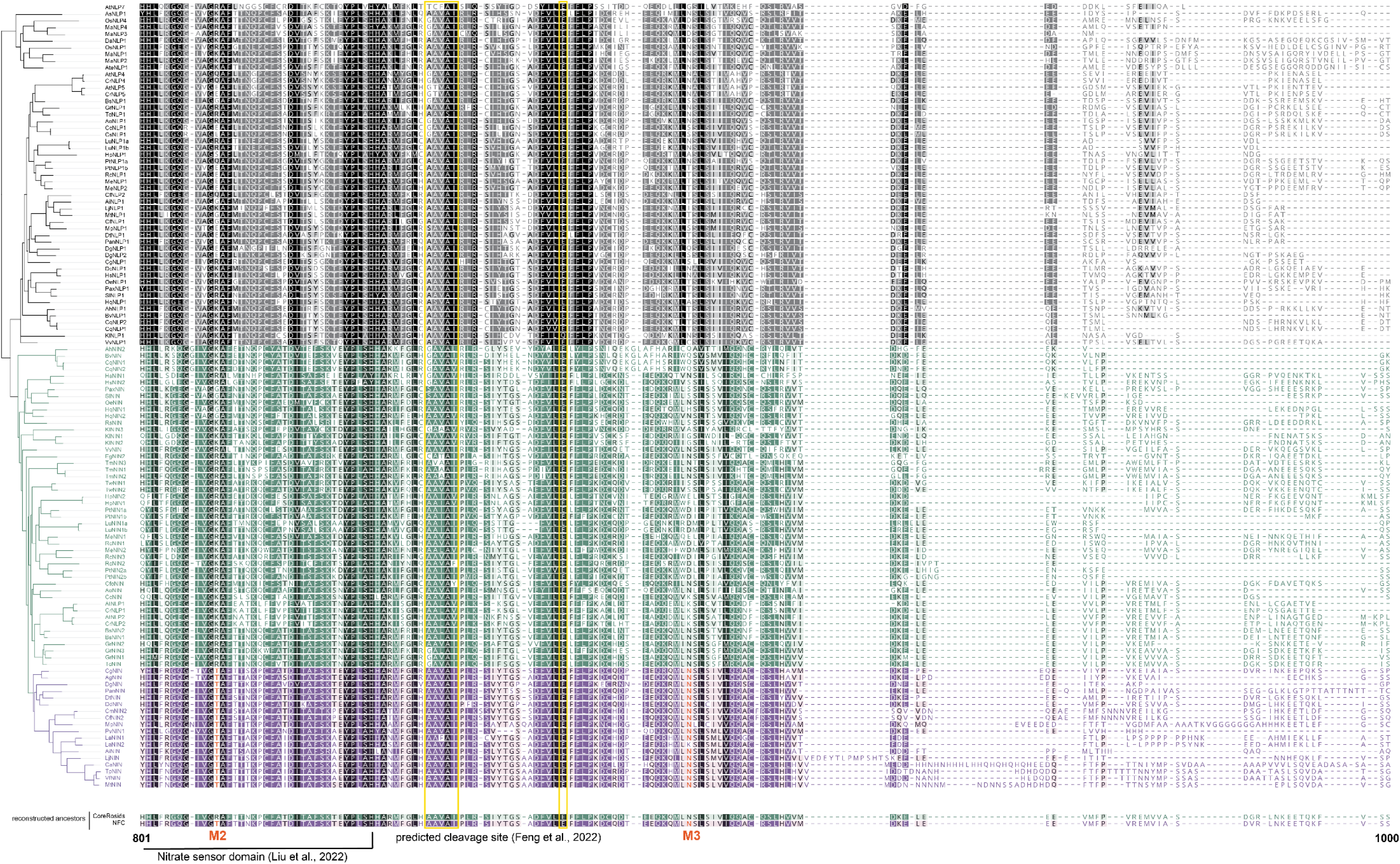
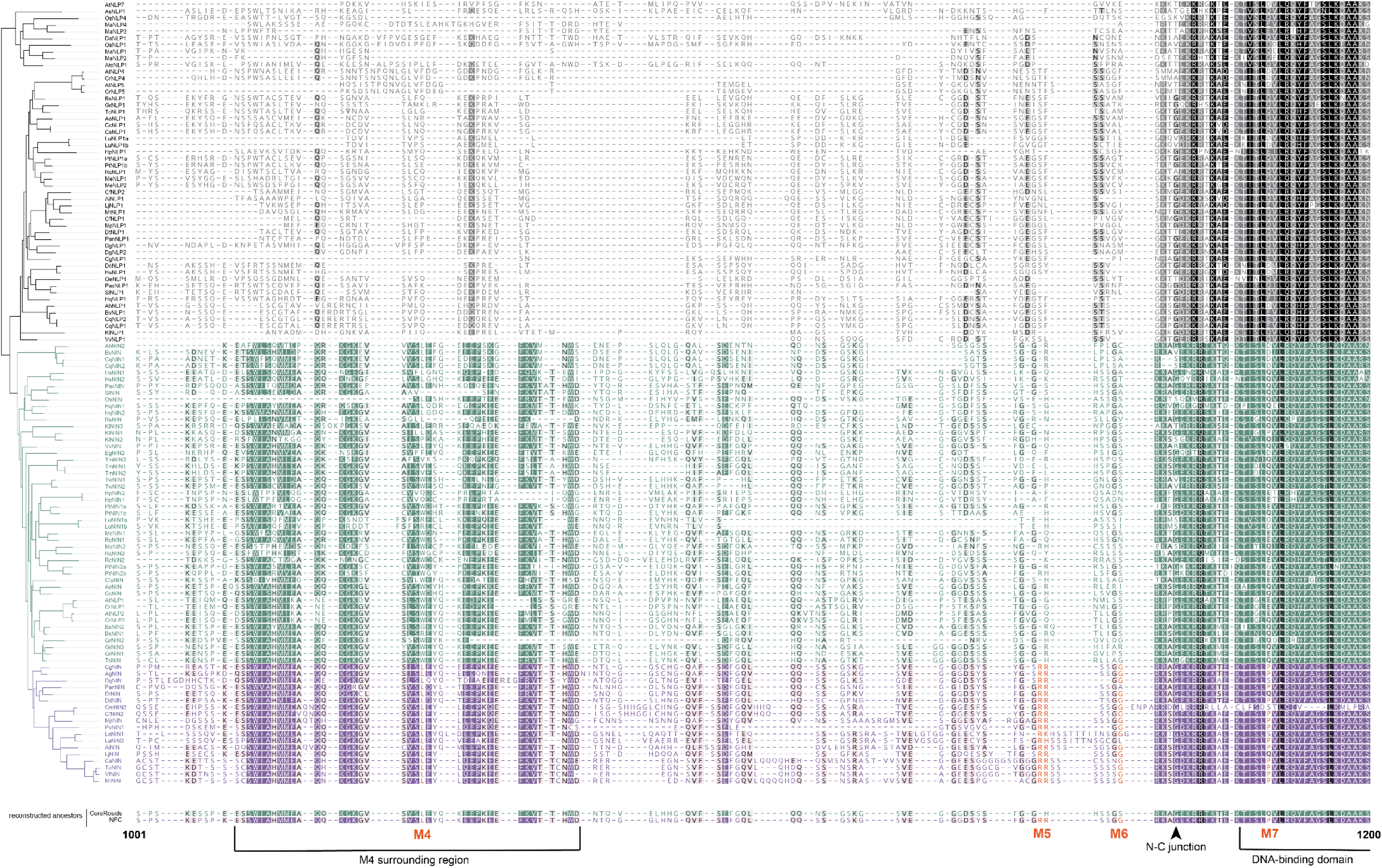
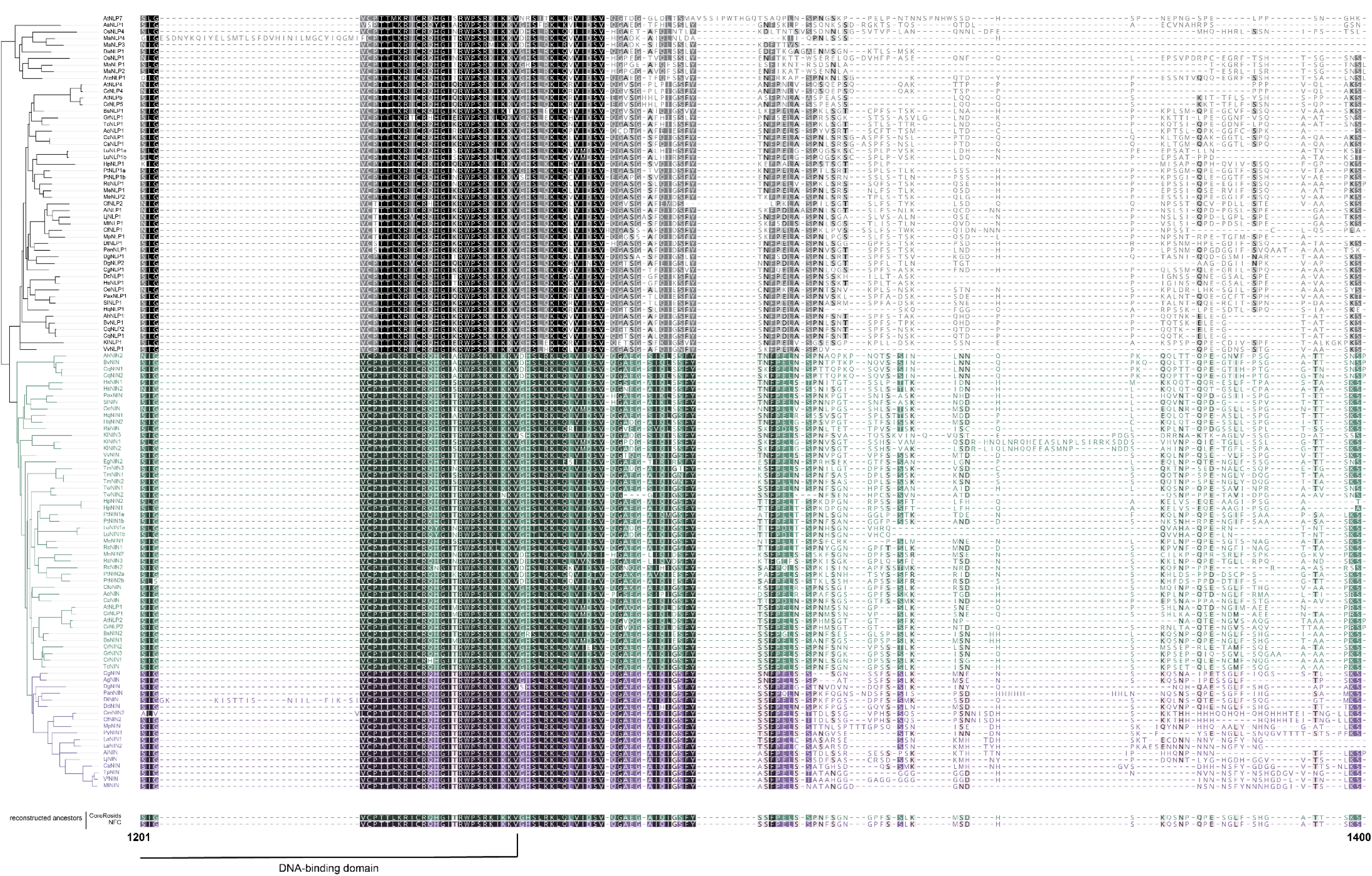
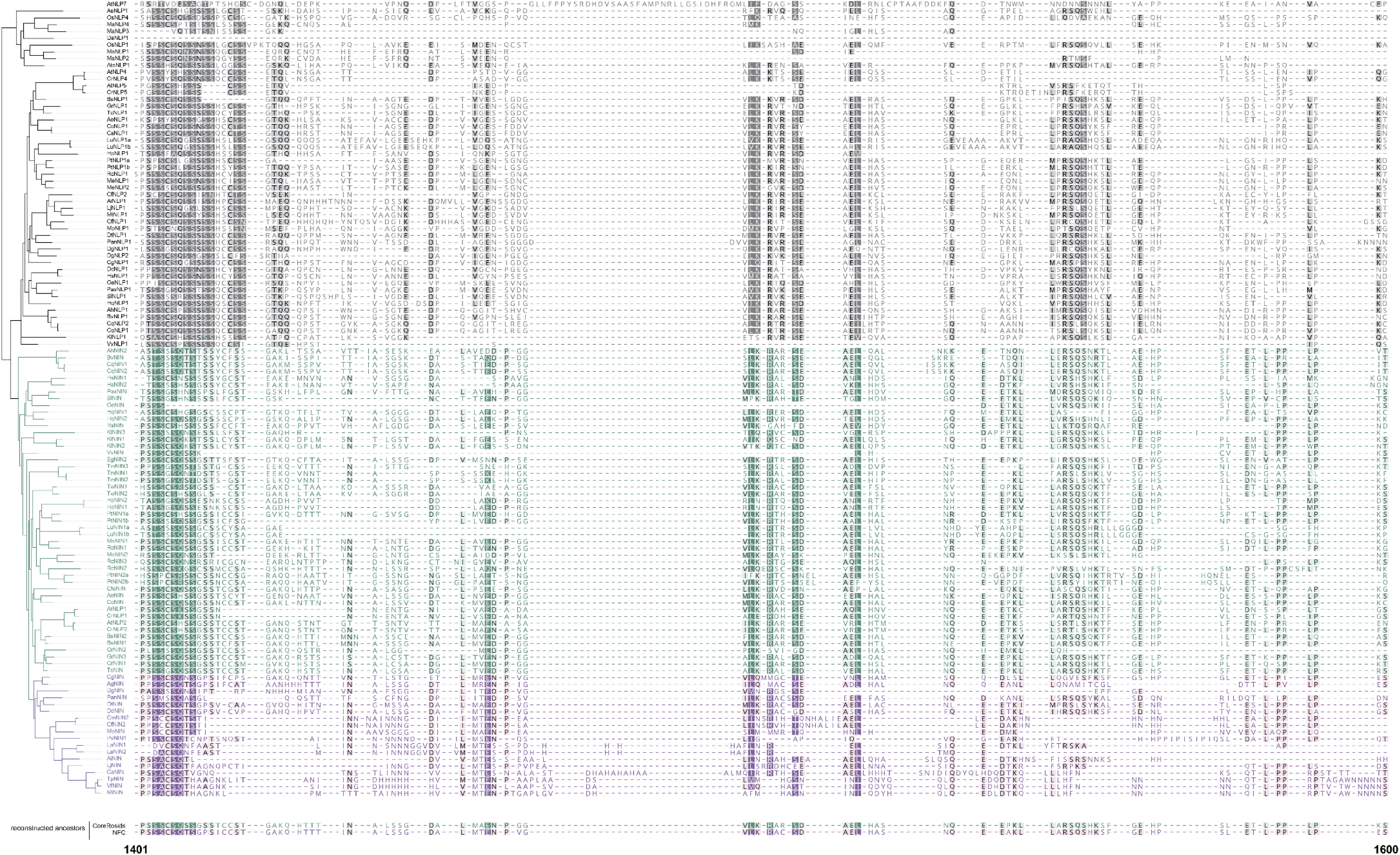
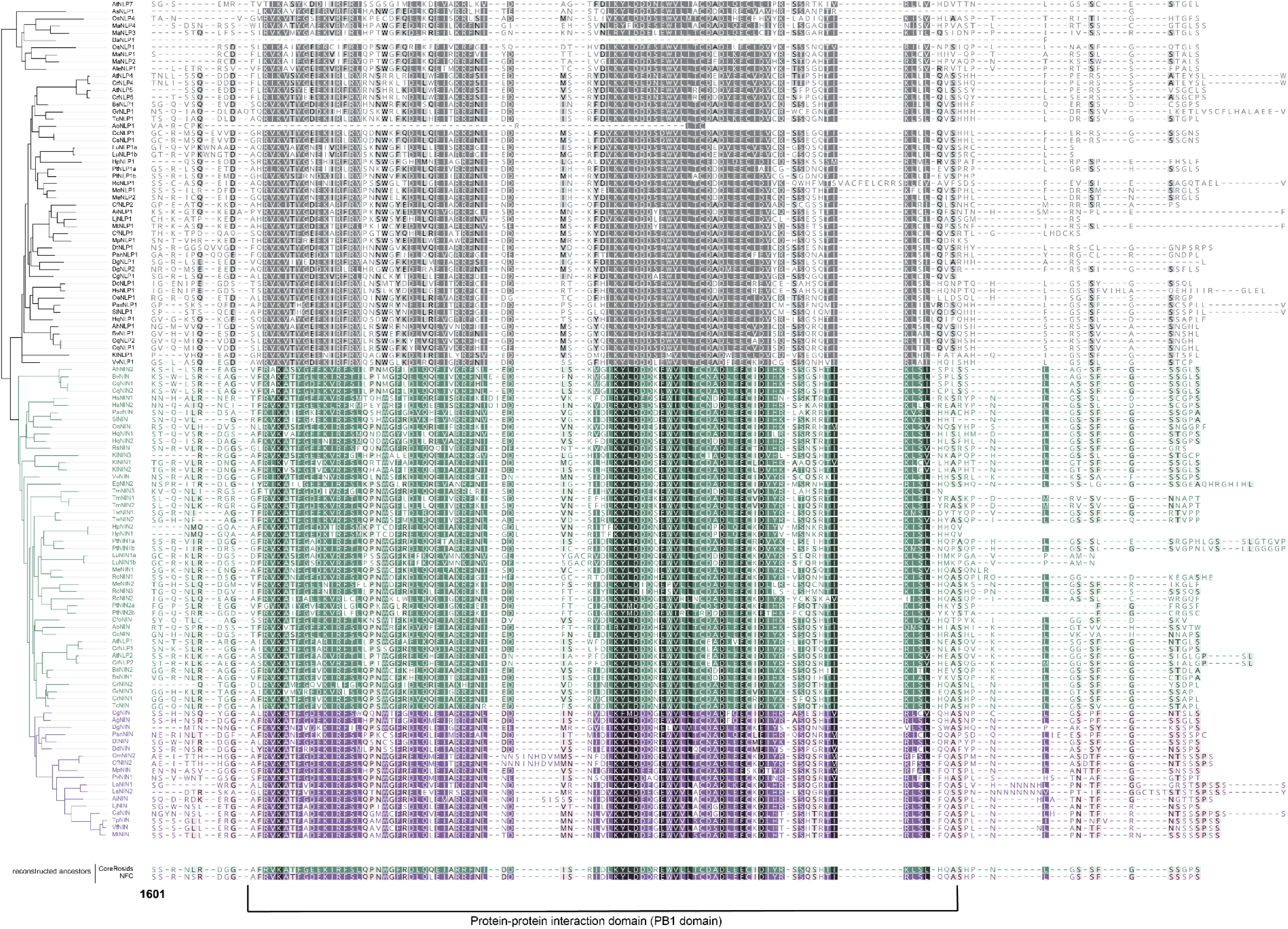
Alignment of NIN and NLP1 orthologs. Multiple sequence alignment of NIN and NLP1 orthologs from a range of angiosperm plant species. Purple (symbiotic NINs) and green (non-symbiotic NINs) shading indicate sequence similarity within the NIN-orthogroup; grey shading indicates sequence similarity within the non-orthologous NLPs. Seven conserved amino acid changes between symbiotic and non-symbiotic NIN orthologs (M1-7) are marked in orange. Numbers below the alignment indicate alignment position including gaps. Ac: *Aquilegia coerulea*, Ag*: Alnus glutinosa*, Ah: *Amaranthus hypochondriacus*, Ai: *Arachis ipaensis,* Ao: *Anacardium occidentale,* As: *Apostasia shenzhenica*, Ate: *Agave tequilana*, At*: Arabidopsis thaliana*, Bs: *Bretschneidera sinensis*, Bv: *Beta vulgaris*, Ca: *Cicer arietinum,* Cc: *Citrus clementina*, Cf: *Chamaecrista fasciculata*, Cfo: *Cephalotus follicularis,* Cg: *Casuarina glauca*, Cm: *Chamaecrista mimosoides*, Cq: *Chenopodium* quinoa, Cr: *Capsella rubella*, Cs: *Citrus sinensis*, Da: *Dioscorea alata*, Dc: *Daucus carota*, Dd*: Dryas drummondii*, Dg: *Datisca glomerata*, Dt: *Discaria trinervis*, Eg: *Eucalyptus grandis*, Gr: *Gossypium raimondii*, Hp: *Hypericum perforatum*, Hq: *Hydrangea quercifolia*, Hs: *Heracleum sosnowskyi*, Kl: *Kalanchoe laxiflora*, La: *Lupinus albus,* Lj: *Lotus japonicus*, Lu: *Linum* usitatissimum, Ma: *Musa acuminata*, Me: *Manihot esculenta*, Mo: *Moringa* oleifera, Mp: *Mimosa pudica*, Mt: *Medicago truncatula*, Oe: *Olea europaea*, Os: *Oryza sativa*, Pan: *Parasponia andersonii*, Pax: *Petunia axillaris*, Pt: *Populus trichocarpa*, Pv: *Phaseolus* vulgaris, Rc: *Ricinus* communis, Rs: *Rhododendron* simsii, Sl: *Solanum lycopersicum*, Tc: *Theobroma cacao*, Tm: *Tetraena* mongolica, Tp: *Trifolium* pratense, Tw: *Tripterygium* wilfordii, Vf: *Vicia* faba, Vv: *Vitis vinifera*, Zm: *Zea mays*. Vertical yellow boxes indicate the previously identified phosphorylation site^30^ and predicted NIN cleavage site^11^. Arrowhead indicates the junction between N- and C-terminus that was used for the constructs in Fig. 2.

**Extended Data Fig. 8.**
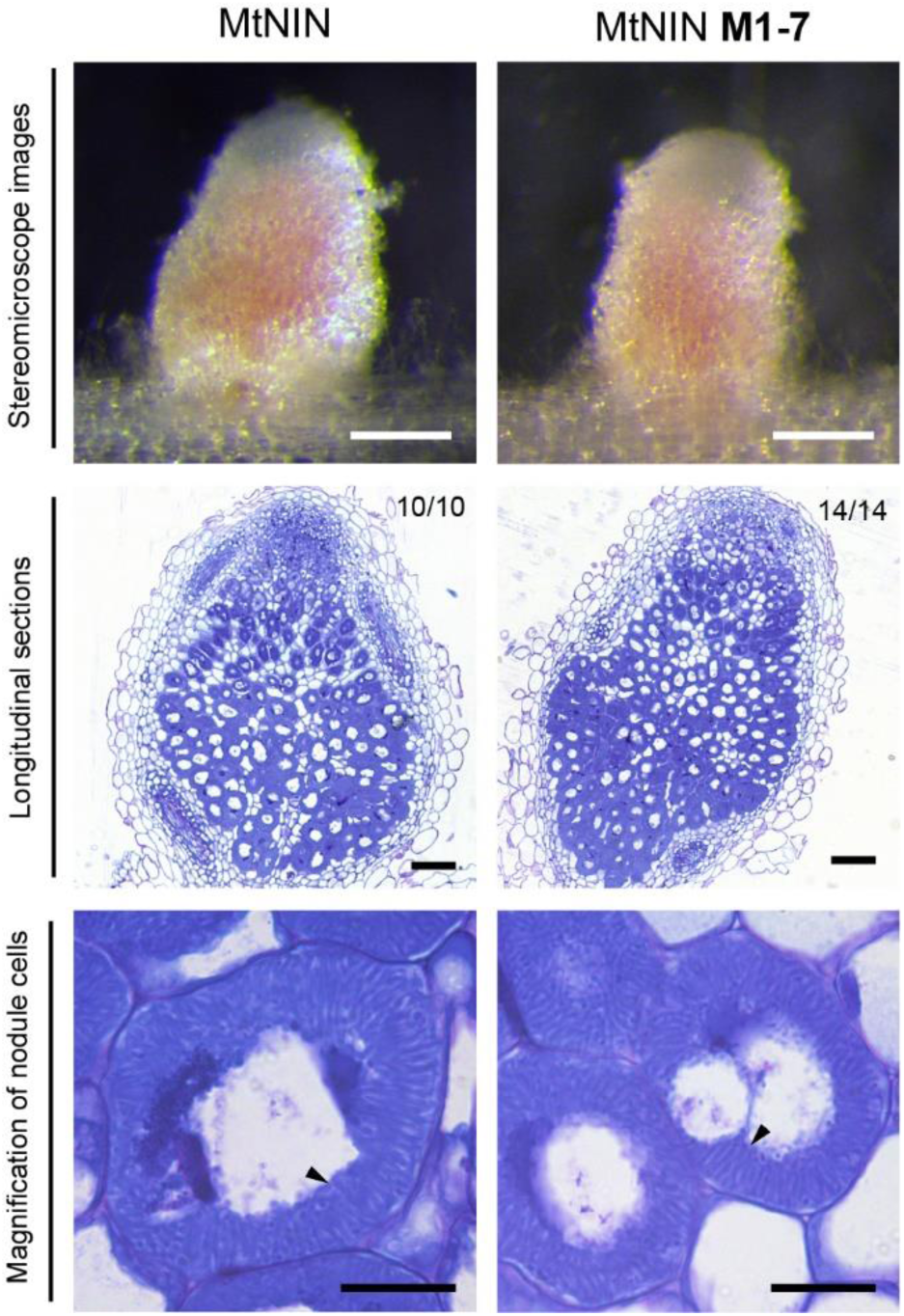
*Mtnin-1* mutant plants complemented by MtNIN^M1–7^ form pink nodules. Nodules formed on *Mtnin-1* mutant roots complemented with MtNIN^M1–7^, in which the seven identified residues (Fig. 4a) of MtNIN have been replaced with their AtNLP2 counterpart. Upper panels: Stereomicroscopic images showing nodules formed on roots complemented with MtNIN^M1–7^ are pink. Scale bars: 2mm. Middle panels: Longitudinal sections stained with toluidine blue. Numbers indicate nodules with released bacteria. Scale bars: 20 μm. Bottom panels: Magnification of nodule cells. Arrowheads indicate fully elongated rhizobia. Scale bars: 20 µm.

**Extended Data Fig. 9.**
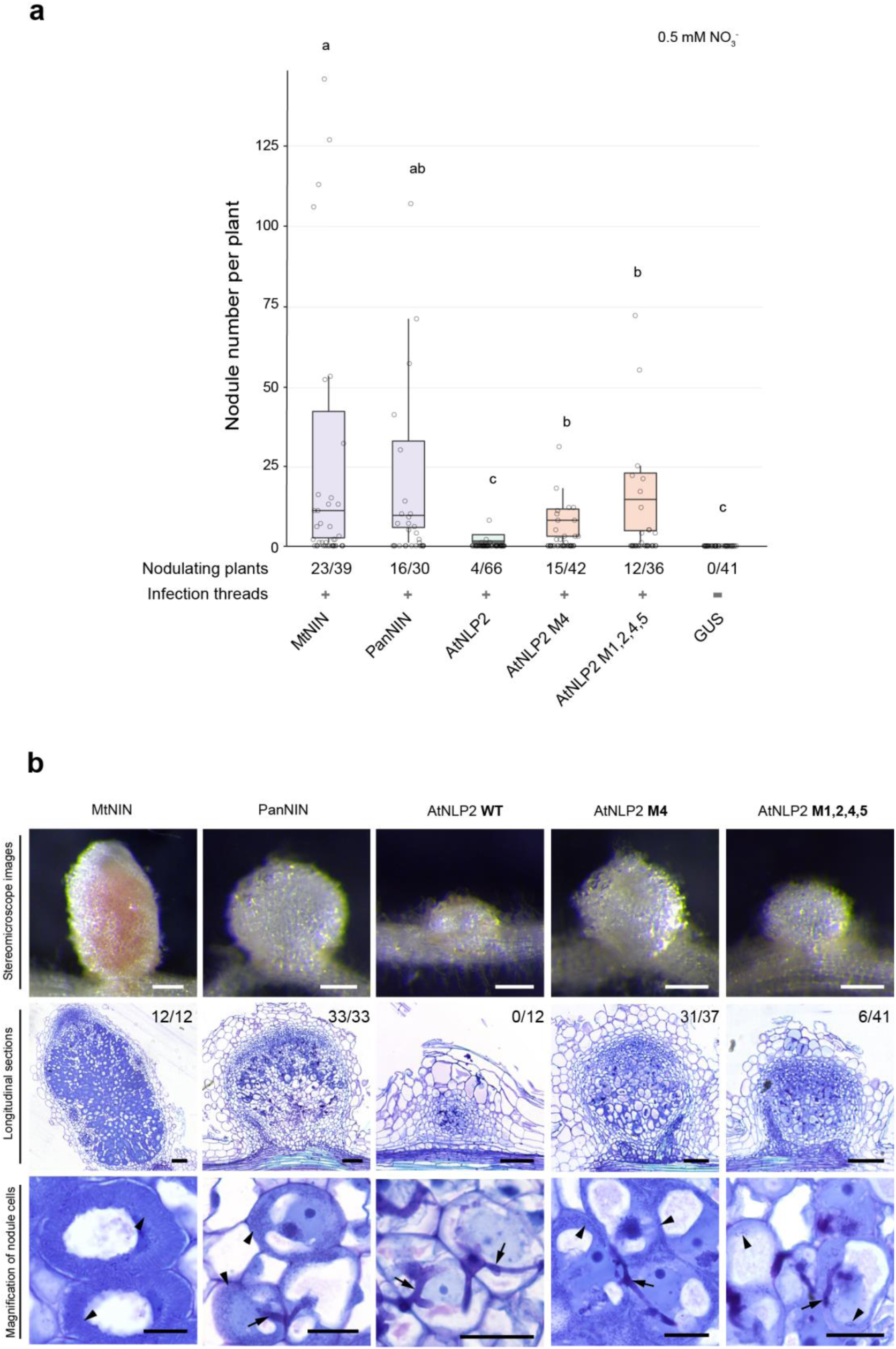
AtNLP2^M4^ functions similarly to symbiotic PanNIN in nodule formation. **a**, Number of nodules formed on *Mtnin-1* mutant roots complemented with MtNIN, PanNIN, AtNLP2, AtNLP2^M4^ and AtNLP2^M1,2,4,5^. Plants were harvested at 4 weeks post inoculation with *S. meliloti* 2011 expressing GFP. Box plots show the number of nodules per nodulated plant. Lowercase letters indicate significant differences between samples (Kruskal-Wallis and post-hoc Dunn’s test, Benjamini-Yekutieli adjusted p < 0.05). **b**, Images of nodules formed on *Mtnin-1* complemented with MtNIN, PanNIN, AtNLP2, AtNLP2^M4^ and AtNLP2^M1,2,4,5^. Upper panels: Stereomicroscope images. Scale bars: 2 mm. Middle panels: Longitudinal sections stained with toluidine blue. Numbers indicate nodules with released bacteria and total nodule numbers studied. Scale bars: 100 μm. Lower panels: Magnification of nodule cells. Arrows indicate infection threads: arrowheads indicate released rhizobia. Scale bars: 20 µm.

**Extended Data Fig. 10.**
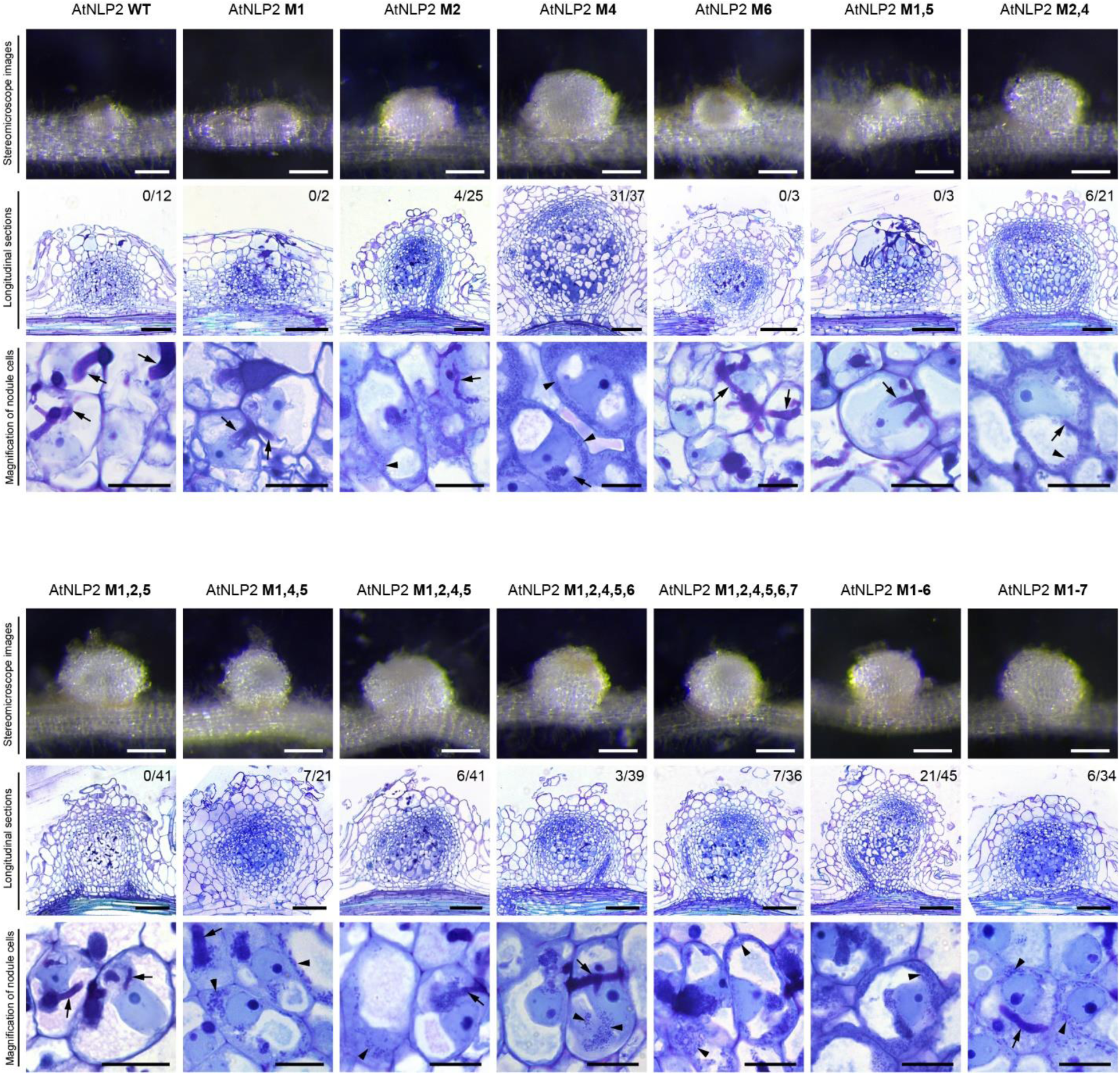
Introducing multiple mutations into AtNLP2 did not further enhance its function in nodule formation relative to AtNLP2^M4^. Images of nodules formed on *Mtnin-1* complemented with AtNLP2 variants, as shown in Fig. 4a,b. The combinations of multiple amino acid substitutions do not further enhance the complementation phenotype relative to the single amino acid substitution AtNLP2^M4^. Upper panels: Stereomicroscope images. Scale bars: 2 mm. Middle panels: Longitudinal sections stained with toluidine blue. Numbers indicate nodules with released bacteria. Scale bars: 100 μm. Lower panels: Magnification of nodule cells. Arrows indicate infection threads; arrowheads indicate released rhizobia. Scale bars: 20 µm.

**Extended Data Fig. 11.**
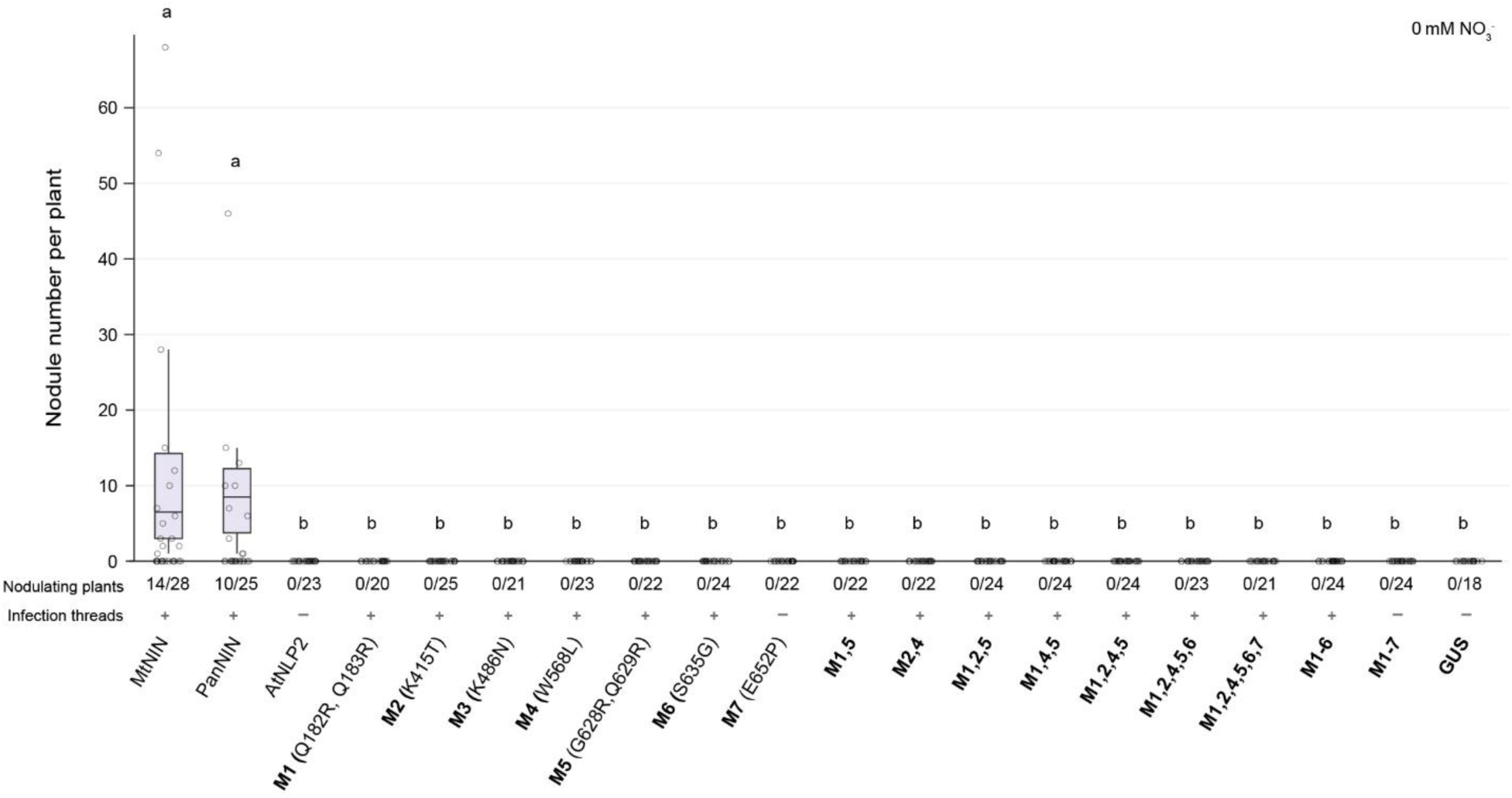
Introducing mutations into AtNLP2 does not improve its symbiotic functionality in absence of exogenous nitrate. Number of nodules formed in absence of exogenous nitrate, on *Mtnin-1* mutant roots complemented with symbiotic NINs, wildtype AtNLP2, and AtNLP2 with different adaptations at single or multiple positions. Lowercase letters indicate significant differences between samples (Kruskal-Wallis and post-hoc Dunn’s test, Benjamini-Yekutieli adjusted p < 0.05).

**Extended Data Fig. 12.**
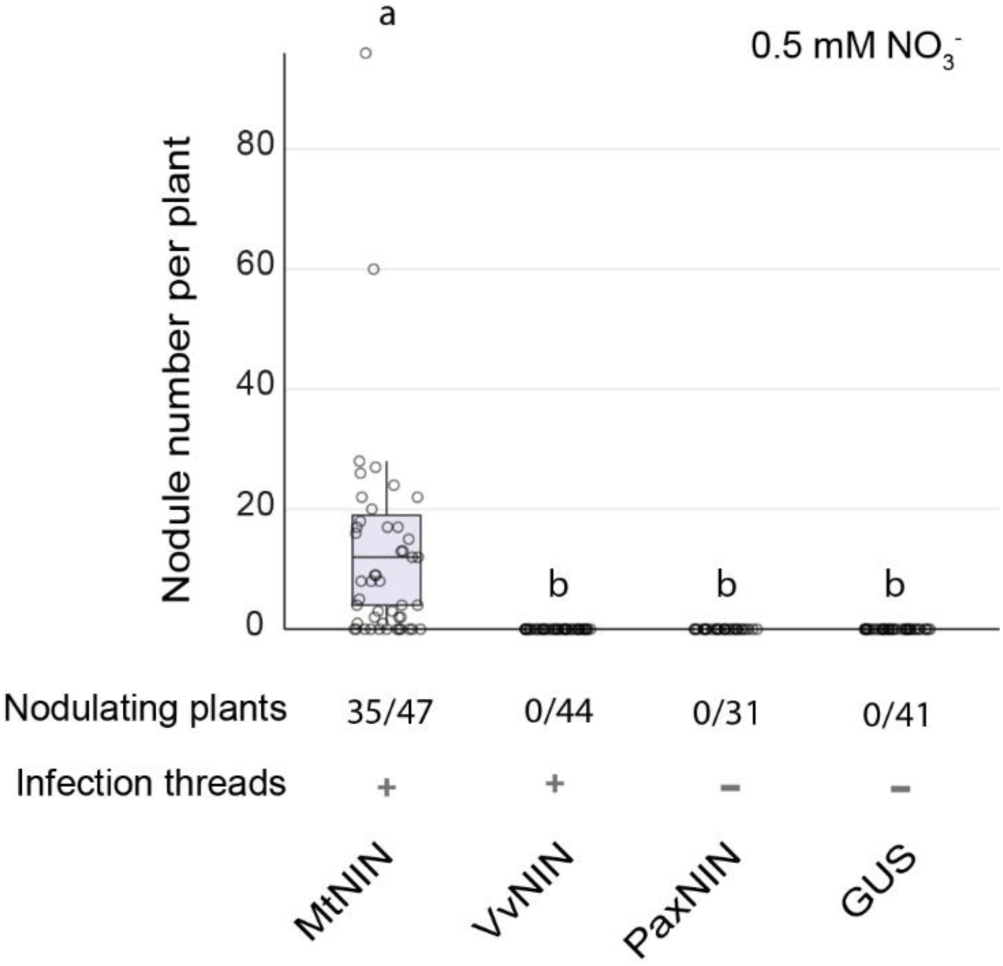
Functionality of Petunia and Vitis NIN in nodule symbiosis. Number of nodules formed on *Mtnin-1* mutant roots complemented with NIN orthologs from *Vitis vinifera* (VvNIN) and *Petunia axillaris* (PaxNIN). Plants were harvested at 4 weeks post inoculation with Sinorhizobium meliloti 2011 expressing GFP. Box plots show the number of nodules per nodulated plant. Lowercase letters indicate significant differences between samples (Kruskal-Wallis and post-hoc Dunn’s test, Benjamini-Yekutieli adjusted p < 0.05).

**Supplementary Table 3.**
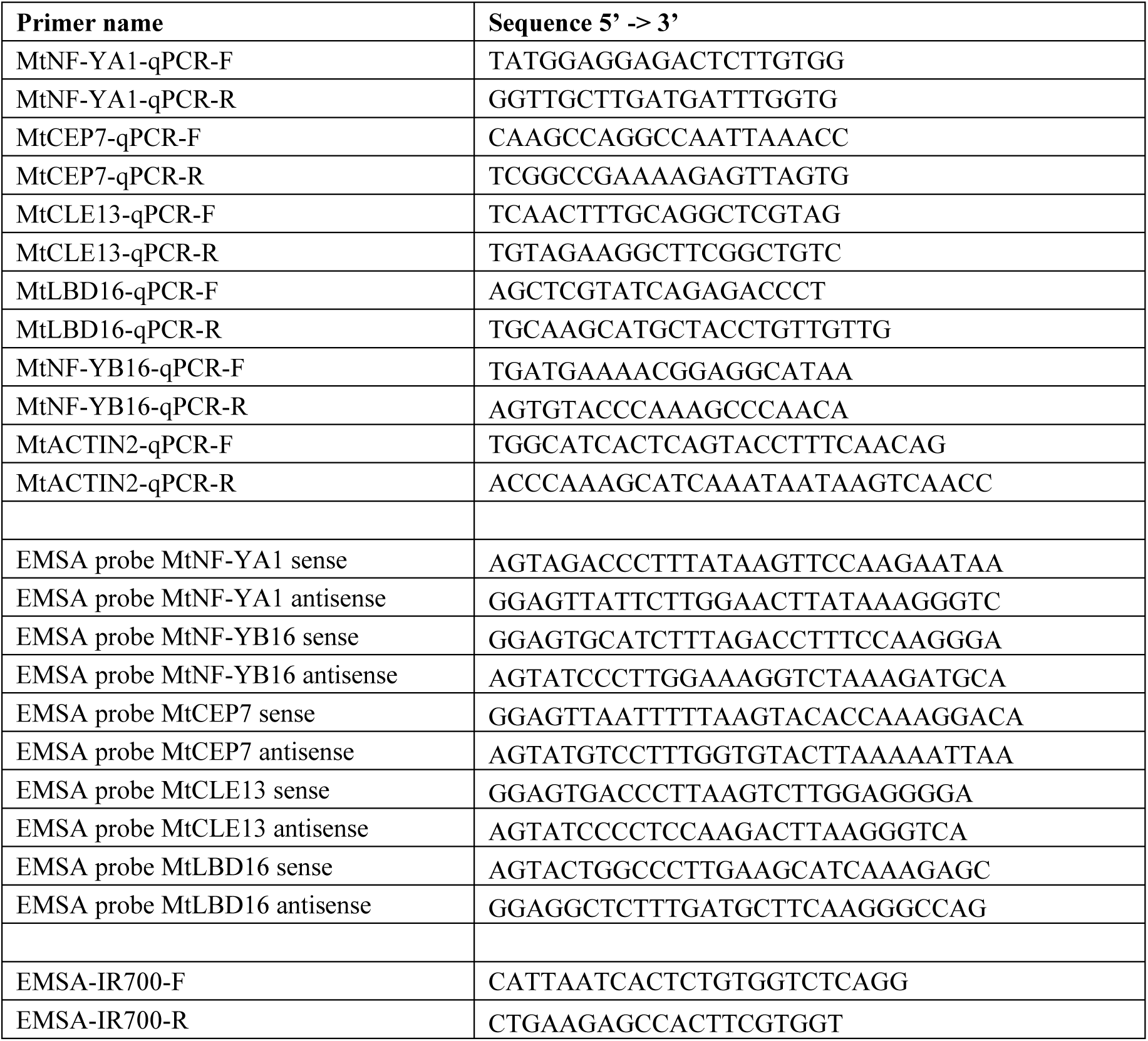
Primers used in this study.

**Supplementary Table 1. (separate file)**

Comparison of codon usage between different NIN orthologs and the Medicago truncatula genome

**Supplementary Table 2. (separate file)**

Ancestral sequence reconstruction and analysis of amino acid conservation

**Supplementary Table 4. (separate file)**

Constructs used in this study

**Supplementary Table 5. (separate file)**

Protein sequences used for analysis

